# Degenerated intervertebral disc environment impairs notochordal cell-derived extracellular vesicles release and their matrix anabolic effect

**DOI:** 10.64898/2026.08.15.744995

**Authors:** Daniele Corraini, Chantal Voskamp, Anouk Eversdijk, Frank M. Riemers, Pieter Vader, Harmjan R. Vos, Keita Ito, Marca H.M. Wauben, Marianna A. Tryfonidou

## Abstract

At the onset of intervertebral disc degeneration, within the disc core, the pH and osmolarity decrease, and the residing notochordal cells (NCs) gradually transition towards nucleus pulposus cells (NPCs). How these microenvironmental cues shape the NC’s extracellular vesicles (EV)-enriched secretome, and thus EV-mediated communication with NPCs during this transition, remains poorly understood. To study this, we collected the secretome from pig NC-rich tissue cultured for 4 days in either healthy or degenerate disc media to mimic these changes. In both conditions, NC-rich tissues were largely comparable at the histological and biochemical levels. Despite, tissues released glycosaminoglycans (GAGs), depleting the extracellular matrix. Surprisingly, degenerative media did not differentially release inflammatory regulators, though it reduced PGE_2_ release. We asked whether this extended to EV-enriched secretome media (SM_EV+), and found that the degenerative media reduced the number of EVs without altering their morphology or size. We then determined NC-EV association of inflammatory and matrix regulators. NC-EV isolation enriched MMP1, IL6 and IL10 and depleted soluble GAGs. Conversely, EV-depletion (SM_EV-) removed most GAGs without affecting MMP1, IL6, and IL10, suggesting that they contribute to the NC-EV soft corona. Functionally, healthy SM_EV+ improved GAG production by NPCs, but attenuated TBXT expression. Degenerate SM_EV+ did not elicit detectable EV-specific effects. These findings suggest that, in health, secretome-mediated communication from NCs to NPCs is only partially EV-mediated. At the onset of IVD degeneration, low pH and osmolarity impair the release of NC-EVs and negate the EV-specific beneficial matrix-anabolic effects on NPCs, contributing to the NC-to-NPC transition.

## Introduction

Low back pain (LBP) is the leading cause of years lived with disability worldwide (Ferreira et al., 2023) and affects over 80% of the human population at least once in their lifetime (O’Sullivan et al., 2019). One of the main causes of LBP is intervertebral disc (IVD) degeneration (Cheung et al., 2009; Luoma et al., 2000). The IVD is responsible for spinal stability and flexibility. Anatomically, it consists of a central gelatinous nucleus pulposus (NP), constrained by the annulus fibrosus, between two cartilaginous endplates that interface with the adjacent bony vertebrae. As the largest avascular organ in the human body, nutrient supply occurs via diffusion, resulting in a progressively limited nutrient availability toward the core of the IVD, the NP (Ren et al., 2023)]. Notably, degeneration of the IVD also starts in its core, the NP, and encompasses multiple processes, with two key aspects being particularly relevant: cellular alterations and changes in the extracellular matrix (ECM), along with shifts in pH and osmolarity (Bartels et al., 1998; Sadowska et al., 2018).

At the cellular level, the NP originates embryonically from the regressing notochord with vacuolated notochordal cells (NCs) residing in the core of the IVD. During disc maturation and the onset of IVD degeneration, the main cell type transitions from NCs to non-vacuolated nucleus pulposus cells (NPCs) (reviewed by Bach et al., 2022). These cellular changes are accompanied by changes in the ECM. In healthy NP the ECM is rich in glycosaminoglycan (GAG) and collagen type II, with a high GAG-to-Collagen type II ratio, which attract water within the NP, leading to an intradiscal osmolarity of 430-496 mOsm (Ishihara et al., 1997; van Dijk et al., 2011), ultimately enabling resistance to compressive loading of the spine column (Risbud et al., 2015; Silagi et al., 2018). During IVD degeneration, this ratio is disrupted by a progress loss and degradation of the GAGs, leading to reduced water content and consequently, lower tissue osmolarity (∼300 mOsm) (Adams & Roughley, 2006; Urban, 2002) alongside a decline in pH from 7.1 towards 6.8 (Bartels et al., 1998). The reduction in pH and osmolarity have been shown to negatively affect the resident NPCs by decreasing cell viability, promoting cell senescence and decreasing cell proliferation and ECM production (Hodson et al., 2018; Laagland et al., 2022; Razaq et al., 2003; Snuggs et al., 2025; Snuggs et al., 2021).

Beyond these structural and cellular changes, IVD degeneration has also been shown to alter the secretome, including all cellular secreted factors of the cells within the NP tissue (Matta et al., 2017). More recently, attention has expanded to the EV-enriched secretome, which refers to the extracellular vesicle (EV)-enriched fraction isolated from conditioned medium, comprising both EVs and co-isolated biomolecules. The EV-enriched secretome of degenerated human NPCs contained more proteins involved in cell-matrix interaction, EV-trafficking, and in stress response, and had lower levels of ECM proteins compared to the secretome of non-degenerated adult human NPCs, suggesting a role for EVs in the pathophysiology of the disc (Li et al., 2024). The specific effect of EVs in the disc remain to be determined, as the bioactive effects of the EV-enriched secretome have been investigated predominantly as a whole, with only a handful of studies specifically examining EV-mediated effects. Furthermore, little is known about the role of the EV-enriched secretome at the onset of IVD degeneration, i.e., during the gradual transition of the residing disc cells from NCs to NPCs. In the context of tissue regeneration, studies have shown that the EV-enriched-secretome of NCs is rich in, ECM-molecules induces cell proliferation and matrix production, and modulates inflammation (Bach et al., 2017; van Maanen et al., 2025). These studies were, however, conducted under supraphysiological disc conditions, i.e., conditioned medium of NC-rich disc tissue was generated in high-glucose and low-osmolarity media, and functional effects were studied on NPCs cultured in high-glucose media too. Furthermore, using EV-depleted methodological controls, it was shown that these NC-secretome effects were only partly EV-mediated (Lan et al., 2019; van Maanen et al., 2025).

Taken together, the influence of the degenerating IVD environment on the NC-secretome and NC-EV characteristics, and their functional effects under IVD-like conditions, warrant further investigation. Hence, the present work focuses on the onset of disc degeneration, studies how environmental changes affect the EV-enriched secretome of NCs and its function in co-residing NPCs, and discerns whether these are EV-mediated. Studies were conducted using the EV-enriched secretome isolated from pig NC-rich tissue. This tissue is rich in NCs and imposes minimal ethical and resource constraints, compared with human NC-rich tissue, which can only be obtained from foetal discs and children (Richardson et al., 2017). Functional studies were conducted in Beagle NPCs, a species that suffers from disc-related disease much like humans do and could serve as a model for both dogs and humans (Thompson et al., 2018).

## Materials and Methods

### Experimental outline

To understand how environmental changes influence NC-EV characteristics and functionality, the EV-enriched secretome was isolated from conditioned medium (NCCM) of pig NC-rich tissues. The tissues were cultured in media that resemble the healthy or degenerate IVD environment, the latter defined by a low pH and osmolarity (**Figure 1**). The effects of these environmental changes on the tissue were evaluated by examining cell morphology, NP marker expression, and ECM composition (histology and biochemical analyses). Furthermore, the NCCM was characterized by (multiplex) immunoassays using a pig-specific panel of inflammatory regulators. The EV secretome, containing NC-EVs and co-isolates, was isolated from NCCM: He-EV and De-EV, respectively. NC-EVs were characterized for morphology (electron microscopy), size (NTA/EM), concentration (NTA), and EV marker expression (proteomic analysis). Lastly, the functionality of EV-enriched secretome, i.e. He-EVs and De-EVs, and corresponding EV-depleted controls was tested in a mild degenerative environment on dog NPC pellets. Pellets were analysed for changes in morphological characteristics, NP marker expression and ECM composition (histology and biochemical analyses).

**Figure 1.**
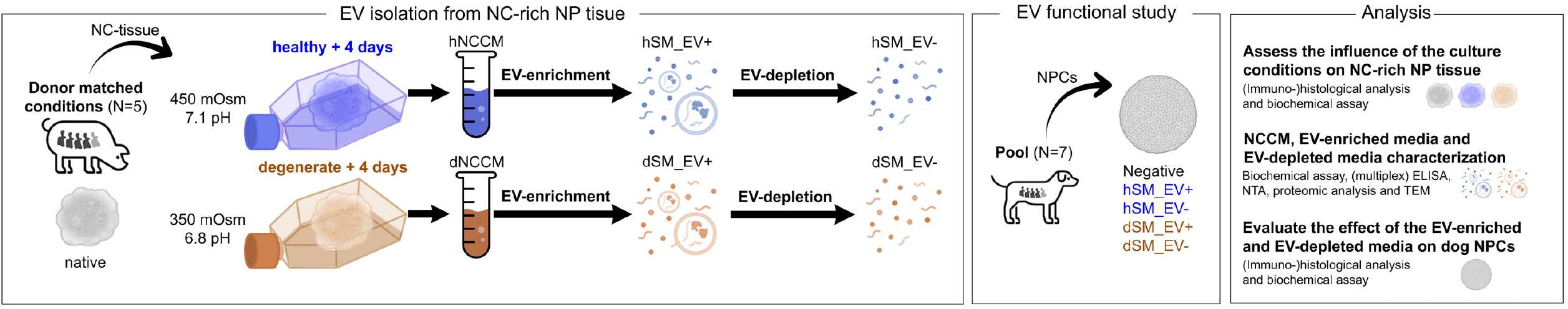
Graphical representation of the experimental workflow. Schematic overview of extracellular vesicle (EV) isolation from notochordal cell (NC) rich nucleus pulposus (NP), functional characterisation and downstream analysis. Healthy environment (h), degenerate environment (d), notochordal cell-conditioned medium (NCCM), EV-enriched secretome medium (SM_EV+), EV-depleted control medium (SM_EV-), nucleus pulposus cells (NPCs), enzyme-linked immunosorbent assay (ELISA), nanoparticle tracking analysis (NTA), transmission electron microscopy (TEM).

### Pig NP tissue collection and conditioned medium production

NP tissues were isolated from complete spines of 5 pigs (*Sus scrofa*, 1.5 months of age, Thompson grade I) supplied by LifeTec Group (Eindhoven), in accordance with Dutch national regulations. The spines were stored at 4°C and NP tissue was collected within 24 hours following sacrifice. For each spine, NP tissue from two lumbar IVDs was collected as baseline. One was stored at −20°C for biochemical analysis and the other was fixed overnight (ON) in 4% paraformaldehyde (PFA) at room temperature (RT) for further histological analysis. The NP tissue, from the remaining IVDs, was used to generate NCCM by culturing NP tissue (1 gram/20 mL medium) either in healthy medium or degenerate medium at 37°C, 5% CO_2_ and 5% O_2_ for 4 days (**Table 1**; Snuggs et al., 2025).

**Table 1:** NP culture media mimicking the healthy and degenerate IVD environment.

|  | Healthy medium | Degenerate medium |
| --- | --- | --- |
| glucose (gr/L) | 1 | 1 |
| pH | 7.1 | 6.8 |
| osmolarity (mOsm/L) | 450 | 350 |

Healthy medium was 1 gr/L low glucose (lg) DMEM (Gibco, 31600083) + 1% GlutaMAX (Gibco, 35050061) supplemented with 11.39 mM NaHCO3 (Sigma-Aldrich, S5761-500G) to set the pH at 7.1, and 94.41 mM NMDG (N-Methyl-D-Glucamine; Sigma-Aldrich, 66930-100G) with 94.41 mM HCl (Sigma-Aldrich, 258148-2.5L) to set the osmolarity to 450 mOsm, without altering the ion-channel potential. In the degenerate medium, pH was set to 6.8 using 5.71 mM NaHCO3, and osmolarity to 350 mOsm using 47.15 mM NMDG+HCl. pH and osmolarity were measured, respectively, with a pH-meter (Mettler Toledo, FiveEasy) and osmometer (Knauer, K-7400S). All media were supplemented with 1% penicillin/streptomycin (P/S; PAA Laboratories, P11-010) and 0.5% fungizone (FZ; Invitrogen, 15290-018).

After 4 days, NCCM was filtered through a 100 μm cell strainer filter, removing the NP tissue. The NP tissue was stored at −20°C for biochemical analysis and PFA-fixed for histological analysis. The filtered NCCM was centrifuged twice at 200 *g* and 500 *g* for 10 minutes at 4°C, to remove tissue debris (**Figure 2**). The NCCM supernatant was collected and an aliquot was stored in low-binding tubes (Thermo Scientific, 90411) at −80°C for biochemical analysis. The NCCM supernatant was 5 times concentrated via centrifugation at 4,000 *g* for 3-6 hours at 4°C using concentration tubes (Pierce™ Protein Concentrators PES Thermo Scientific™, Pierce™ Protein Concentrator PES, Volume=20 mL, MWCO=3K, Thermo Scientific, 88526, 24/pack). Subsequently, the concentrated NCCM was centrifuged at 10,000 *g* for 40 minutes at 4°C to remove cellular debris and apoptotic bodies. The supernatant was aliquoted into low-binding tubes and stored at −80°C until EV isolation.

**Figure 2.**
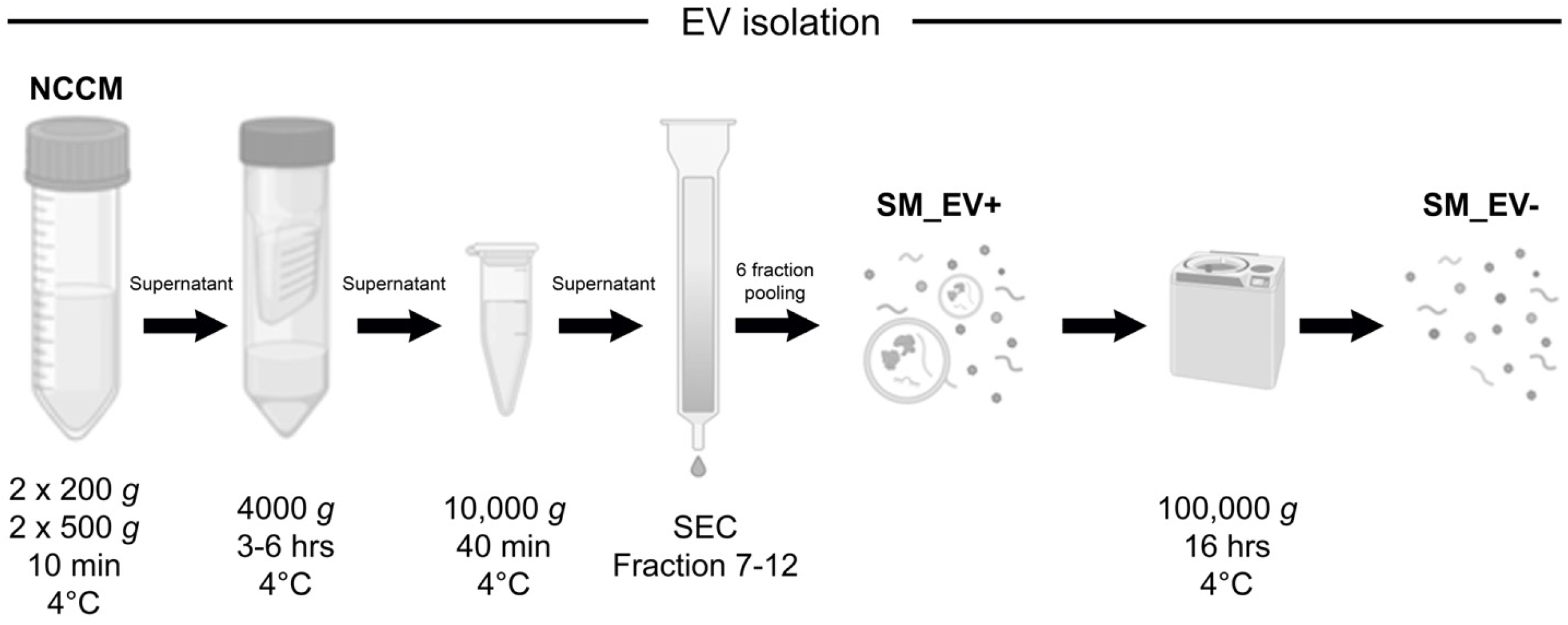
Isolation of extracellular vesicle (EV)-enriched secretome medium and EV-depleted control medium from notochordal cells. Schematic overview of EV isolation using differential centrifugation followed by size exclusion chromatography. Notochordal cell-conditioned medium (NCCM), EV-enriched secretome medium (SM_EV+), EV-depleted control medium (SM_EV-).

### EVs isolation

NC-EVs were enriched from NCCM within 6 months via Size-Exclusion Chromatography (SEC) using qEV SEC-Columns (iZON Science, qEVoriginal Legacy, 70nm, 500 µL), loading 1 mL samples/column as previously reported (**Figure 2**) (Bach et al., 2017; van Maanen et al., 2023). The qEV columns were washed with degenerate medium containing 1% P/S + 0.5% FZ. Per qEV column, 1 mL of 5x concentrated NCCM was added and the samples were eluted with degenerate medium containing 1% P/S + 0.5% FZ. 15 fractions of 0.5 mL were collected per qEV column, and their respective protein concentrations were determined (DeNovix, DS-11, A280). Based on previous EV characterization (van Maanen et al., 2023), 6 fractions were collected and pooled, yielding 3 mL per culture condition (**Supplementary figure 1A**). This process was repeated twice, yielding 6 mL of EV-enriched secretome medium from NCCM generated under healthy (hSM_EV+) and degenerate (dSM_EV+) conditions.

To the eluted SM_EV+ media (6 mL), growth factor-free discogenic supplements for NPC culture were added to prepare degenerate disc medium, at final concentrations of 1% ITS-X (Thermo Scientific, 51500056), 0.04 mg/mL L-proline (Sigma, P5607), and 1.25 mg/mL human serum albumin (HSA, CSL Behring GmbH, 15526054). SM_EV+ media were aliquoted into low-binding tubes and stored at −80°C until further use. To generate respective EV-depleted media (SM_EV-), 4 mL of SM_EV+ media for each condition were transferred into SW60 tubes and ultracentrifuged (UC) at 100,000 g for 16 hours at 4°C (SW60 rotor; 32,100 rpm; RCF average 10,016 g; RCF max 13,648 *g*; κ-factor 2543.1). The 100,000g UC pellet was discarded, and the UC supernatant of each condition was aliquoted in low-binding tubes and stored at −80°C until further use. All media were supplemented with fresh 0.1 mM ascorbic acid 2-phosphate (ASAP; Sigma, A8960) before being added to the NPC pellets.

### (Multiplex)immunoassays of NCCM and EV media inflammatory regulators content

The effects of the healthy and degenerate culture conditions on NP tissue at the inflammatory level were studied by measuring PGE2 and CCL2 levels in NCCM using enzyme-linked immunosorbent assay (ELISA) kits (PGE_2_ – 514010, Cayman chemical; CCL2 - PB0088S-100, PBB0089S-050, Kingfisher Biotech) following the manufacturer’s instructions. The results were acquired with a microplate reader (BMG Labtech, CLARIOstar). Undiluted NCCM was assessed for secreted inflammatory regulators, including TNF, IL1B, IFNG, IL1RN, IL6, IL10, CXCL8 and MMP1 with a pig-specific magnetic bead-based multiplex assay (LXSAPM, Biotechne) according to the manufacturer’s protocol. Results were acquired with a Luminex analyser (MAGPIX®, Luminex Corporation). Furthermore, the levels of these inflammatory regulators were determined for Healthy and Degenerate SM_EV+ and SM_EV− samples.

### NC-EV characterization

#### Nanoparticle tracking analysis

EV size distribution and concentration profiles were inferred from measurements obtained by nanoparticle tracking analysis (NTA)for both SM_EV+ and SM_EV− media, for each testing condition and pig spine, and compared to the negative controls represented by plain degenerate medium and PBS used to dilute the samples before the NTA analysis (1:1000). NTA was performed with NanoSight NS500 (Malvern Panalytical) using 405 nm laser, sCMOS camera and NTA 3.4 software. Each sample was diluted with Dulbecco’s PBS (Sigma-Aldrich, D8537) to achieve 40-150 particles per frame. For each sample, five 30-second captures were acquired, with a temperature set at 25°C and viscosity 0.9 cP. The camera level was set to 16, with a shutter speed of 1300, a gain of 512, a frame rate of 25 frames per second, at detection threshold of 20, and automatic blur size.

#### Transmission electron microscopy

NC-EV morphology was determined via transmission electron microscopy (TEM). NC-EVs were isolated as previously described and eluted in PBS, then concentrated 10 times via centrifugation at 4000*g* for 3 hours at 4°C using concentration tubes (Pierce™ Protein Concentrators PES Thermo Scientific™, Pierce™ Protein Concentrator PES, MWCO=3K, Thermo Scientific, 88526). Briefly, for TEM NC-EV samples were bound on fresh carbon-coated Formvar copper grids (hexagonal mesh 200). The unbound sample was washed away with PBS, and the bound sample was fixed with 1% glutaraldehyde in PBS. Then the grids were rinsed extensively with MQ-water and stained with 0.5% uranyl oxalate, pH 7.0, for 10 min at RT. Thereafter, they were transferred into ice-cooled Uranyl-Methylcellulose pH 4.0 for 10 min at RT. Excess Uranyl-Methylcellulose was removed by blotting, and grids were air-dried for at least 1 hour at RT. Samples were imaged with a JEOL3 electron microscope equipped with a VELETA SO4U fast CCD sensor and 4-megapixel camera at 80 kV.

#### Proteomic analysis

The presence and abundance of common EV marker (Welsh et al., 2024) were determined in both healthy and degenerate SM_EV+ and SM_EV− media via proteomics analysis. The samples were dried in the SpeedVac after which 40 µl 150mM HEPES (pH 8), 2% SDS, 10 mM Tris(2-carboxyethyl)phosphine hydrochloride (TCEP), 40 mM 2-chloro-acetamide (CAA) was added to denature the proteins and alkylate the cysteines. Proteins were cleaned up with the SP3 protocol (Hughes et al., 2014), using 10 µL beads (500 µg). Proteins were digested overnight at 37 °C on a heater-shaker with 0.5 µg Trypsin/LysC (Thermo) in 100 mM HEPES (pH 8), 10 mM CaCl_2_. After acidification with 2% Formic Acid (FA), peptides were separated from the beads and were loaded on C-18 stage tips (Affinisep). After elution from the stage tips, acetonitrile was removed using a SpeedVac and the remaining peptide solution was diluted with buffer A (0.1% FA) before loading. Peptides were separated after trapping on a pre-column on a 25 cm in-house made pulled emitter fused silica column (50 µm ID, Polymicro) packed with 1.9 µm aquapur gold C-18 material (dr. Maisch) using a 1-hour gradient (7% to 80% ACN 0.1% FA), delivered by a Vanquish Neo HPLC (Thermo), and electro-sprayed directly into an Orbitrap Astral Mass Spectrometer (Thermo Scientific). The latter was set in data independent mode with a cycle time of 0.6 seconds for both Faims compensation voltage (CV) settings (−45V and −65V), in which the full scan was performed at a resolution of 240K. The precursor mass range for both CVs was 380-980 Th, and the peptides, isolated within a 2 Th window, were fragmented at a normalized collision energy of 25% and an AGC of 500%.

The raw mass spectrometry data were searched using Thermo Scientific Proteome Discoverer Software (version 3.2.0.450) with the Chimerys DIA workflow. The reference proteome of Sus Scrofa (Pig) organism, downloaded from UniProt in December 2021, was used for protein identification; a set of common contaminant sequences was also included. Enzymatic digestion was specified as trypsin/P, allowing up to two missed cleavages. Carbamidomethylation was set as a fixed modification, while methionine oxidation was treated as variable modification. The maximum mass tolerance for fragment peaks was set to 5 ppm. Protein quantification was based on MS2 Apex signal intensities, and a false discovery rate threshold of 1% (q-value ≤ 0.01) was applied at both the peptide and protein levels. Protein group abundance intensities were extracted from the report and used for further analysis. The proteomics data have been deposited to the ProteomeXchange Consortium via the PRIDE partner repository (http://www.ebi.ac.uk/pride) with the dataset identifiers.

The protein group output was further processed in R (version 4.3.3) for downstream analysis. Proteins that were identified underwent pre-processing, retaining only those with at least 2 unique peptides and identified in at least 3 out of 5 replicates per condition. Average protein abundance across biological replicates was background-corrected and normalized through a variance-stabilizing transformation (vsn). Protein isotypes were aggregated and averaged. Investigated EV markers belonged to 4 different classes: transmembrane proteins, cytosolic proteins, co-isolated proteins and non-endosomal intracellular proteins. EV markers of interest that were not detected in any condition were left out of the graphical display.

### EV functional study on dog NPC pellets under degenerate discogenic condition

#### NPC isolation and expansion

NPCs were isolated from NP tissue of complete spines from 7 dogs with mild to moderate disc degeneration (*Canis familiars*, beagle, male, 19-43 months old, Thompson grade II-III, **Table 2**). Beagles were terminated in unrelated research studies, overseen by the Local Welfare Body and approved by the National Authorities for Animal Experiments. NP tissue was collected within 24 hours after euthanasia. To isolate NPCs, NP tissue was digested with 5.25 U/mL pronase (Roche Diagnostics, 11459643001) at 37°C for 30 minutes and with 62.5 U/mL collagenase type II (Worthington, 4176) at 37°C for 4-16 hours. Isolated NPCs were expanded in hgDMEM (Invitrogen, 31966), 450 mOsm (52.5 mM NMDG) with 10 % fetal bovine serum (FBS, Gibco 10500-064, lot number 08F0597K, Thermo Fisher Scientific), 1 % P/S, 0.5% FZ, 0.1 mM ASAP in 5% O_2_, 5% CO_2_ at 37 °C. After 4-7 days, NPCs were trypsinised at sub-confluency and reseeded at a density of 6,000 cells/cm^2^. NPCs in passage 2 were used for experiments.

**Table 2:** Information overview of Beagle dogs used to isolate NPCs presenting with Thompson grade II-III.

| Dog | Sex | Age (months) |
| --- | --- | --- |
| A | male | 19 |
| B | male | 19 |
| C | male | 22 |
| D | male | 22 |
| E | male | 31 |
| F | male | 42 |
| G | male | 43 |

#### Pellet formation and NC-EV functional study

To assess the donor-specific effect of the NC-EVs on a representative dog NPC population and a relevant *in vitro* 3D NPC model, NPCs from seven male dogs were pooled in equal numbers before forming pellets. Pellets were formed by plating 35,000 NPCs per well in a ultra-low attachment 96-wells plates (Costar, 7007) in 50 µl of hgDMEM + Glutamax, 450 mOsm (52.5 mM NMDG+HCl) supplemented with 1% P/S, 0.5% fungizone, 0.1 mM ASAP and discogenic factors (1.25 mg/mL HSA, 1% ITS-X, and 0.04 mg/mL L-proline) plus 10 ng/mL recombinant human TGF-β1 (*h*TGF-β1 R&D Systems, 240-B-010), to induce matrix deposition. Pellet formation was induced by centrifuging the 96-well plates at 500 rpm for 5 minutes. The pellets were cultured in 5% O_2_, 5% CO_2_ at 37 °C for 3 days. After 3 days, media were refreshed to same media but without *h*TGF-β1 (i.e. 50 µL of fresh hgDMEM + Glutamax supplemented with discogenic factors, 1% P/S, 0.5% fungizone, 0.1 mM ASAP) for 1day, whereafter three pellets were collected and processed for further biochemical and histological analysis to determine baseline levels. On day 4, the medium was replaced with 50 µL of low glucose medium, to mimic the low-glucose IVD environment (**Supplementary figure 2A**). NC-EV functionality was studied in degenerate medium supplemented with discogenic factors, 1% P/S, 0,5% FZ and fresh 0.1 mM ASAP using the following conditions, i.e. 1) Control; 2) hSM_EV+; 3) hSM_EV−; 4) dSM_EV+; 5) dSM_dEV− media.

For all experiments the EV-depleted control, either hSM_EV− or dSM_EV−, represents the methodological controls for the EV samples. A positive control in healthy medium supplemented with 10 ng/mL *h*TGF-β1 was included. The NPC pellets were cultured for 14 days in 5% O_2_, 5% CO_2_ at 37 °C. Media were refreshed twice a week, collected and stored at −20°C. After 14 days of treatment, pellets were imaged with the stereomicroscope (EVOS FL Imaging System, AMF4300) for pellet size determination using ImageJ software (version 1.54t). Pellet size was quantified as the pellet projected area by applying a threshold mask for the dark area on the brightfield background. Pellets were then collected and washed twice with Hank’s Balanced Salt Solution (HBSS) to remove culture medium before being processed for further biochemical and histological analysis. To confirm NC-EV-specific effects, follow-up studies were conducted by serially diluting the He- and De-EV media 1:10 and 1:100 in degenerate medium using the same culture conditions as previously described. The anabolic effects of NC-EVs on NPC pellets were assessed at 14 days of culture using GAG and DNA content.

### NP tissue and pellet histological analysis

Histological analyses was performed on pig NP tissues and dog NPC pellets. They were fixed in 4% PFA ON at RT, followed by dehydration processing and embedding in paraffin. To analyse the cellularity and cell morphology, 5 µm sections were stained with haematoxylin and eosin (H&E) (Merck Millipore, 1.09249.2500). The average cross-sectional area of NC vacuoles present in pig NP tissue was quantified on H&E-stained sections via ImageJ software (version 1.54t). A threshold mask of the unstained area was applied to identify the individual intracellular regions, each representing a vacuolated structure. The cross-sectional area of each intracellular region was measured and averaged per section to determine the average cross-sectional area of NC vacuoles.

To analyse the quality of the deposited matrix, sections were stained with toluidine blue (TB; Sigma-Aldrich, C.I. 42040) following standard protocols. To detect NP tissue-specific cellular proteins, immunohistochemical staining for T-box transcription factor T (TBXT) and cytokeratin 8+18+19 (KRT8/18/19) was done. For ECM components, aggrecan (ACAN), collagen type 1 (COL1) and collagen type 2 (COL2) were evaluated. To further characterize the NPC model employed in the EV functional studies, dog NPC pellets were also tested for expression of the NP marker angiopoietin-1 receptor (TIE2), which is commonly expressed by NPC progenitors. Furthermore, expression of TIE2 is specifically associated with the production of a healthy ECM (Guerrero et al., 2021; Sakai et al., 2018).

For immunohistochemical staining, sections were rehydrated (xylene and 100%, 96%, 70% ethanol), blocked with 0.3% H_2_O_2_ (Sigma, H1009) for 10 min and washed twice with 0.1% PBS/Tween-20 (PBS-T). Thereafter, depending on the antibody, antigen retrieval was performed (**Supplementary table 1**). Sections were washed twice with PBS-T and blocked accordingly (**Supplementary table 1**). Primary antibody was added and incubated ON at 4°C. As a negative control, the appropriate isotype was used at the same concentration as the respective primary antibody. After incubation, sections were washed twice in PBS-T and the respective secondary antibody was added for 30 min at RT. Sections were then washed with PBS and incubated with 3,3’-diaminobenzidine solution (Bright-DAB, VWR, BS04-110) for 7 min. The chromogenic reaction was stopped with one wash in demineralized water. Sections were counterstained with haematoxylin QS solution (Sigma-Aldrich, MHS32) for 1 min and washed in running tap water for 10 min. Stained sections were dehydrated (70%, 96%, 100% ethanol, and xylene) and mounted. Stained sections were imaged with BX-43 microscope (CellSens Imaging software, Olympus) and immunopositivity was quantified as a percentage of positive stained area via ImageJ software (version 1.54t). Images were subjected to ImageJ Colour Deconvolution to separate DAB positive staining from H/E and automatic ImageJ threshold was applied to DAB staining images. Percentage of positive area of DAB staining images was determined by manually selecting the complete sample section area and automatically measuring the amount of positive stained area over the total selected sample section area. TBXT immunopositivity was also quantified as a percentage of positively stained nuclei using QuPath-0.6.0 (version 0.6.0). Images were subjected to automatic QuPath DAB-H/E Colour Deconvolution and automatic QuPath threshold was applied to DAB staining images. Percentage of positively DAB stained nuclei was automatically determined measuring the amount of positive nuclei over total amount of nuclei automatically identified by QuPath software.

### DNA and GAG biochemical analysis of NP tissue and pellet and media

Biochemical analysis was conducted to quantify (a) the effect of the two culture environments on the DNA and GAG content of the pig NP tissue, and (b) the functional effects of the He-EV and De-EV on DNA and GAG content of the dog NPC pellets. To determine their respective wet and dry weights, NP tissues were weighed, dehydrated (Speedvac System, Savant), and weighed again. The dehydrated NP tissues and NPC pellets were digested, respectively in 750 µL and 50 µL, of papain digestion solution (10 mM papain; Sigma, P3125-100mg), 1.57 mg/mL cysteine HCl (Sigma, C9768-5G) at 60°C ON. The next day, samples were vortexed, subjected to 1 more hour at 60°C in the papain digestion solution, and stored at −20°C. The DNA concentration of the digested NP tissues and pellets was measured using the 1x high sensitivity (HS) dsDNA Qubit kit (Invitrogen, 15860210) according to the manufacturer’s instructions. The GAG content of the digested NP tissues, He- and De-NCCM, He-EV and De-EV media and respective EV-depleted controls, digested dog NPC pellets, and pellet culture media was measured via 1,9-dimethylmethylene blue (DMMB) assay (Farndale et al., 1982). Upon addition of DMMB (Sigma, 341088), absorbance at 525 and 595 nm was measured via a microplate reader (BMG Labtech, CLARIOstar). GAGs concentration was determined using a chondroitin sulfate (Sigma, C4384) standard curve. Cumulative GAG released from the dog NPC pellets was defined as GAGs released in the media over time, corrected for the GAG amount of the respective EV or EV-depleted media: Cumulative GAG released = ∑(t)(pellet culture medium GAG amount – EV medium GAG amount).

### Statistical analysis

Statistical analyses were performed with GraphPad Prism 10.4.1. First, data were tested for normal and log normal distribution with the Shapiro-Wilk test, and otherwise confirmed with Q-Q plot due to the small sample size. To control for inter-donor variability, analyses were performed using donor-matched samples. If data were normally distributed, a one-way analysis of variance (ANOVA) was used assuming data sphericity for paired data and applying Šidák’s multiple comparison correction. If the data were not normally distributed, the nonparametric Friedman’s test was used with Dunn’s multiple-comparison correction. Statistical significance of differences between the negative condition and the He-EV or De-EV conditions was determined via an unpaired t-test with Welch’s correction. A *p*-value < 0.05 was considered statistically significant. Given the large inter-donor biological variability, a *p*-value between 0.05 and 0.1 was considered as a trend towards statistical significance. Statistically insignificant comparisons are omitted in graphs.

## Results

### Effect of degenerative environment on NC phenotype and matrix

To understand the effect of the degenerative environment on NCs embedded within the ECM, the NC phenotype and NC-matrix were analysed after 4 days in degenerative culture conditions and compared to healthy conditions. The DNA content of the tissue remained stable across both environments, indicating that the cell number was not affected. (**Figure 3C).** Furthermore, NCs retained their characteristic vacuoles (**Figure 3A**). Quantification of the cross-sectional area of these vacuoles on H&E-stained histological sections revealed an increased area upon tissue culture in a healthy (2-fold increase) and degenerative environment (1.3-fold), with the latter not reaching statistical significance (*p*=0.4; **Figure 3B**). Besides, immunopositivity of TBXT (**Figure 3D-3E)** and KRT8/18/19 **(Figure 3G-3H**) was maintained under degenerate and healthy culture conditions. Nuclear immunopositivity for TBXT, indicative of active TBXT signalling, was increased in both healthy and degenerative culture conditions, however without reaching statistical significance due to the high inter-donor variability (respectively *p*=0.3 and *p*=0.9; **Figure 3F**).

**Figure 3.**
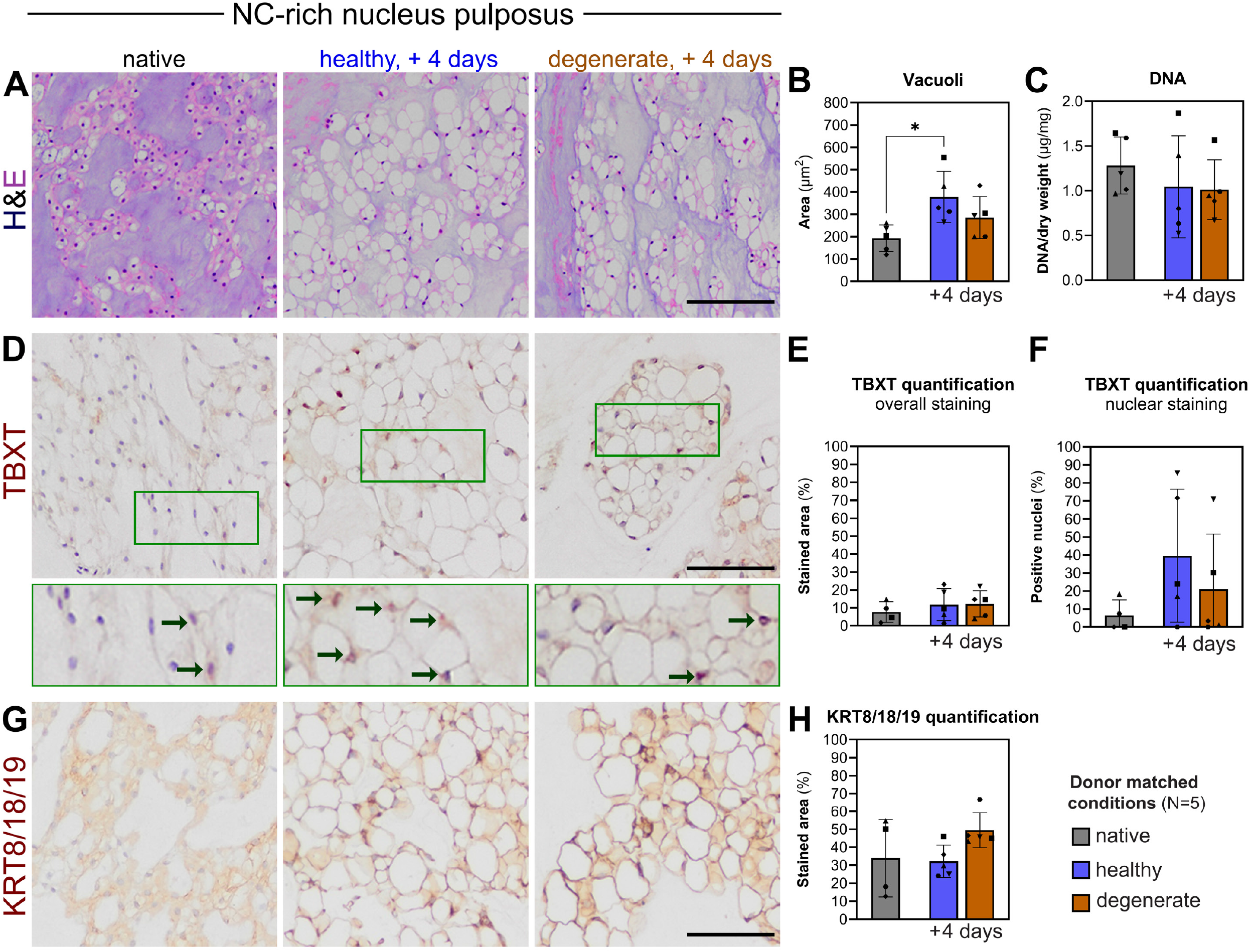
Effect of healthy and degenerate medium on pig notochordal cell (NC) phenotype. All analyses were done on native porcine nucleus pulposus (NP) tissues or tissues that were cultured in either healthy or degenerate medium for 4 days. (**A-B**) Haematoxylin and Eosin (H&E) staining of NP tissues (scale bar represents 200 µm) with corresponding quantification of NC vacuoli area. (**C**) DNA content of NP tissues normalized by dry weight. (**D-F**) Immunohistochemistry for brachyury (TBXT) with corresponding quantification of the percentage of positive stained area and positive stained nuclei (scale bar represents 200 µm). (**G-H**) Immunohistochemistry for cytokeratin 8/18/19 (KRT8/18/19) with corresponding quantification of the percentage of positive stained area (scale bar represents 100 µm). Dots indicate individual matched pig donor (N=5). Bars and whiskers indicate means ± SD. *p*-values are indicated for statistical significance of differences between conditions using Sidak’s multiple comparison test (* : *p* < 0.05).

Next, we determined the effect of the environment on the ECM. Toluidine blue staining was detected in all tissues, but was less intense in tissues cultured in degenerative or healthy environments compared with native tissue (**Figure 4A**). In line with this, GAG tissue content decreased to comparable levels upon culture under both healthy and degenerate conditions (*p=*0.6), by ∼80% and 75% respectively (**Figure 4B**). Accordingly, GAG was released into the medium at comparable levels under both culture environments (*p*=0.2; **Figure 4C**). This observation prompted us to measure the osmolarity and determine whether the released GAGs altered the osmotic environment. Notably, the osmolarity of He-NCCM and De-NCCM upon 4 days of tissue culture was comparable to the respective osmolarity set at day 0 (**Supplementary figure 3E**). Furthermore, ACAN immunopositivity decreased in healthy (*p*=0.05; trend) and degenerate culture conditions (**Figure 4D-4E**), while COL2 immunopositivity levels were maintained (**Figure 4F-4G**), and COL1 immunopositivity levels remained undetectable (**Figure 4H-4I**).

**Figure 4.**
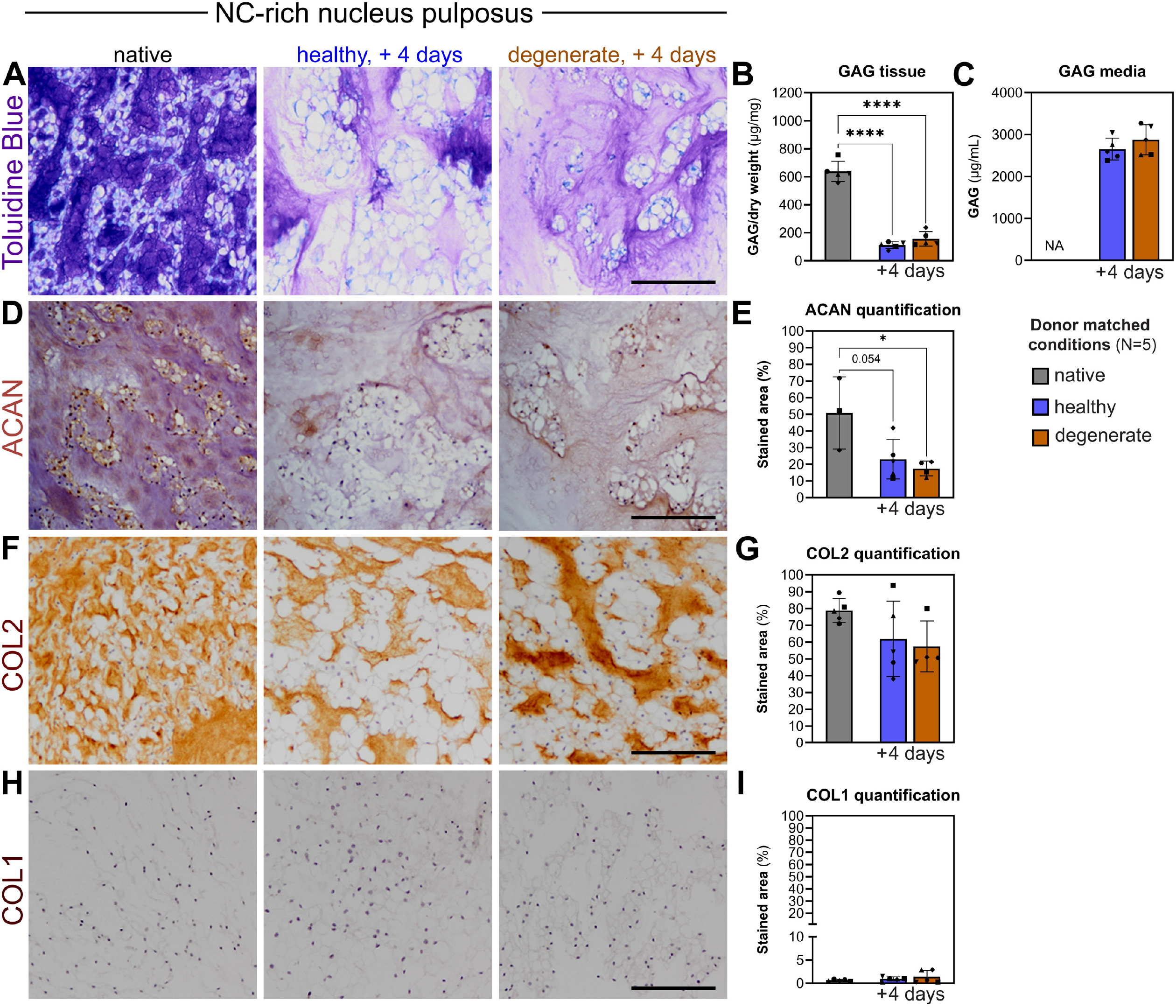
Effect of healthy and degenerate medium on pig notochordal cell (NC) extracellular matrix. All analyses were done on native porcine nucleus pulposus (NP) tissues or tissues that were cultured in either healthy or degenerate medium for 4 days. (**A**) Toluidine Blue staining (scale bar represents 200 µm). (**B**) Glycosaminoglycans (GAG) content in tissues normalized by dry weight. (**C**) GAG released from the NP tissues upon culturing – paired t-test analysis. (**D-I**) Immunohistochemistry for aggrecan (ACAN), collagen 2(COL2) and collagen 1 (COL1) with corresponding quantification of the percentage of positive stained area (scale bar represents 200 µm). Dots indicate individual matched pig donor (N=5). Bars and whiskers indicate means ± SD. *p*-values are indicated for statistical significance of differences between conditions using Sidak’s multiple comparison test (numeric *p*-value: *p* < 0.1, * : *p* < 0.05, ** : *p* < 0.005, *** : *p* < 0.0005, **** : *p* < 0.0001). *p* between 0.05 – 0.1 is considered a statistical trend and provided in absolute values.

Altogether, the data indicate that in this experimental setup, the NC phenotype and matrix characteristics in the healthy and degenerative environments were comparable at the histological and biochemical level. Free-swelling tissue culture resulted in GAG depletion in both conditions and reduced confinement of the NCs, leading to larger vacuoles. In the healthy medium, vacuoles were further enlarged, likely due to the higher osmolarity of the medium (Hunter et al., 2007).

### Degenerative environment affects inflammatory mediator PGE2 levels in NCCM

Next, we investigated the effect of the culture environment on the release of inflammatory mediators from the NC-rich tissue by measuring their levels in NCCM. CCL2 and IL10 were released in dNCCM and hNCCM at comparable levels (**Figure 5A-5C**). Moreover, TNF, IL1B, IFNG, IL6, IL1RN, CXCL8, and MMP1 levels in the NCCM were under the detection limit in both conditions (**Supplementary figure 3A-3D**). Surprisingly, lower PGE2 media levels were measured in the dNCCM than in the hNCCM (**Figure 5D**). Overall, NCs embedded in their matrix unexpectedly did not increase the release of inflammatory regulators and even reduced PGE2 release under a mildly degenerative culture environment (350 mOsm and 6.8 pH).

**Figure 5.**
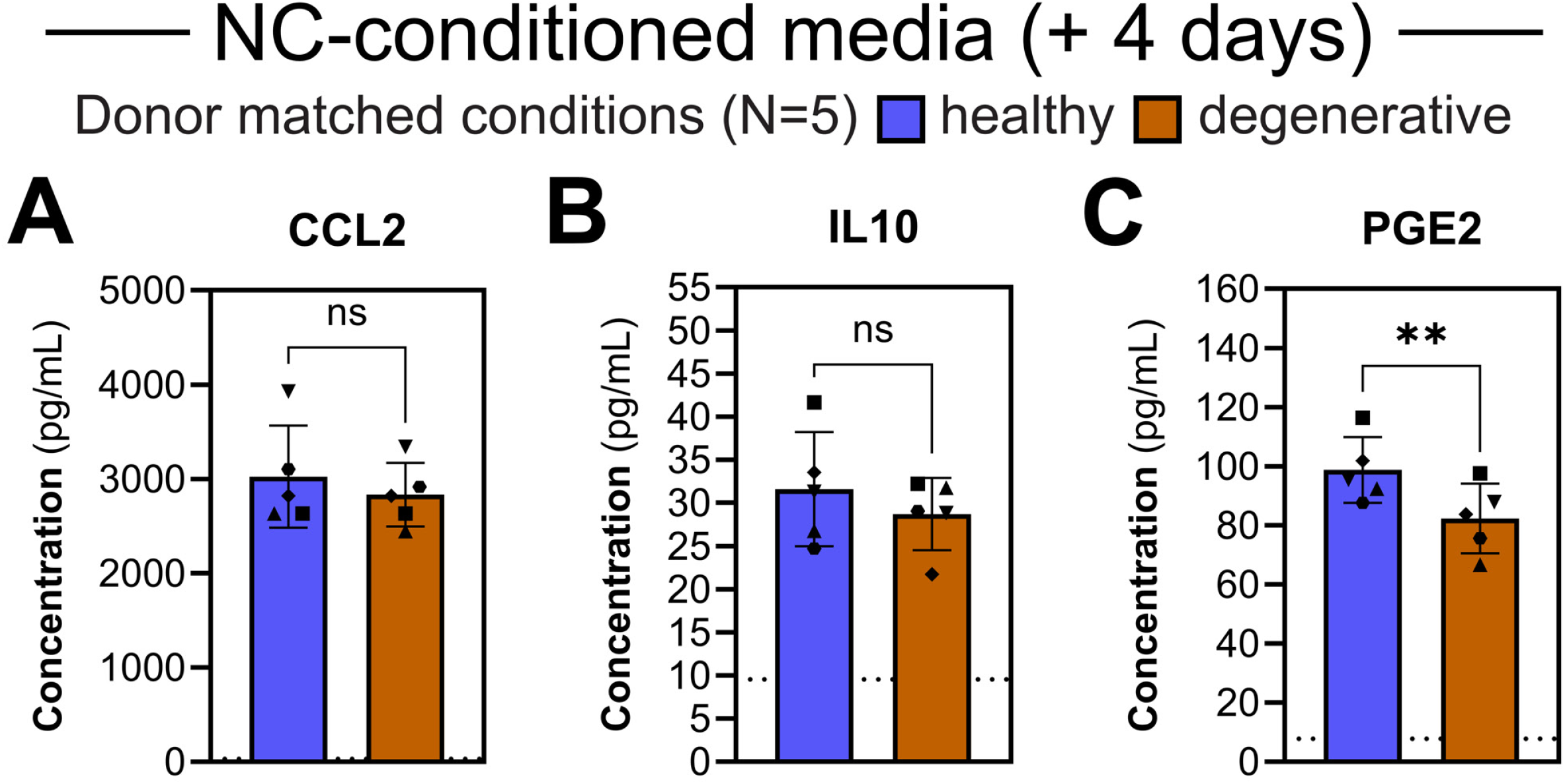
Inflammatory profile of notochordal cell conditioned medium (NCCM) following culture in healthy or degenerate media. All analyses were done on NCCM derived from porcine nucleus pulposus (NP) tissue cultured in healthy or degenerate medium for 4 days. (**A-C**) Levels of CCL2, IL10 and PGE2 in healthy or degenerate NCCM. Dotted line represents the Multiplex ELISA detection limit of the respective analyte (CCL2: 31.3 pg/mL; IL10: 9.5 pg/mL; PGE2: 7.8 pg/mL). Dots indicate individual matched pig donor (N=5). Bars and whiskers indicate means ± SD. *p*-values are indicated for statistical significance of differences between conditions using paired Student t-test analysis (ns : not significant, **: *p* < 0.005).

### A degenerative environment decreases the number of released NC-EVs

Next, we investigated the effect of the degenerative environment on the release of NC-EVs and their characteristics, including size distribution, morphology and the presence of EV markers through proteomic analysis. Nanoparticle tracking analysis showed that the dSM_EV+ had a particle size distribution comparable to hSM_EV+, spanning 100 to 350 nm, with a peak around 200 nm (**Figure 6A**) and their morphology was confirmed by transmission electron microscopy (**Figure 6C**). Notably, dSM_EV+ particle concentration was two times lower than that of hSM_EV+ (**Figure 6B**). The respective EV-depleted procedural controls were also analysed. The average particle concentration confirmed EV depletion by 99% for both hSM_EV− and dSM_EV− (*p*=0.0003 and *p*=0.06, respectively; **Figure 6B**). Moreover, the particle size distribution of hSM_EV− and dSM_EV− spanned 50 to 300 nm, and was comparable between conditions (**Figure 6A; Supplementary figure 1B**). At the protein level, the abundance of EV markers, including CD9, CD63 and TSG101, was higher in both hSM_EV+ and dSM_EV+ compared to the respective EV-depleted samples (**Figure 6D**). Hence, the EV isolation protocol successfully enriched for NC-EVs and the EV-depletion protocol was effective in depleting NC-EVs. Altogether, these results show that cues from a degenerative environment lowered EV release from NC-rich tissue, while preserving their morphology and size distribution.

**Figure 6.**
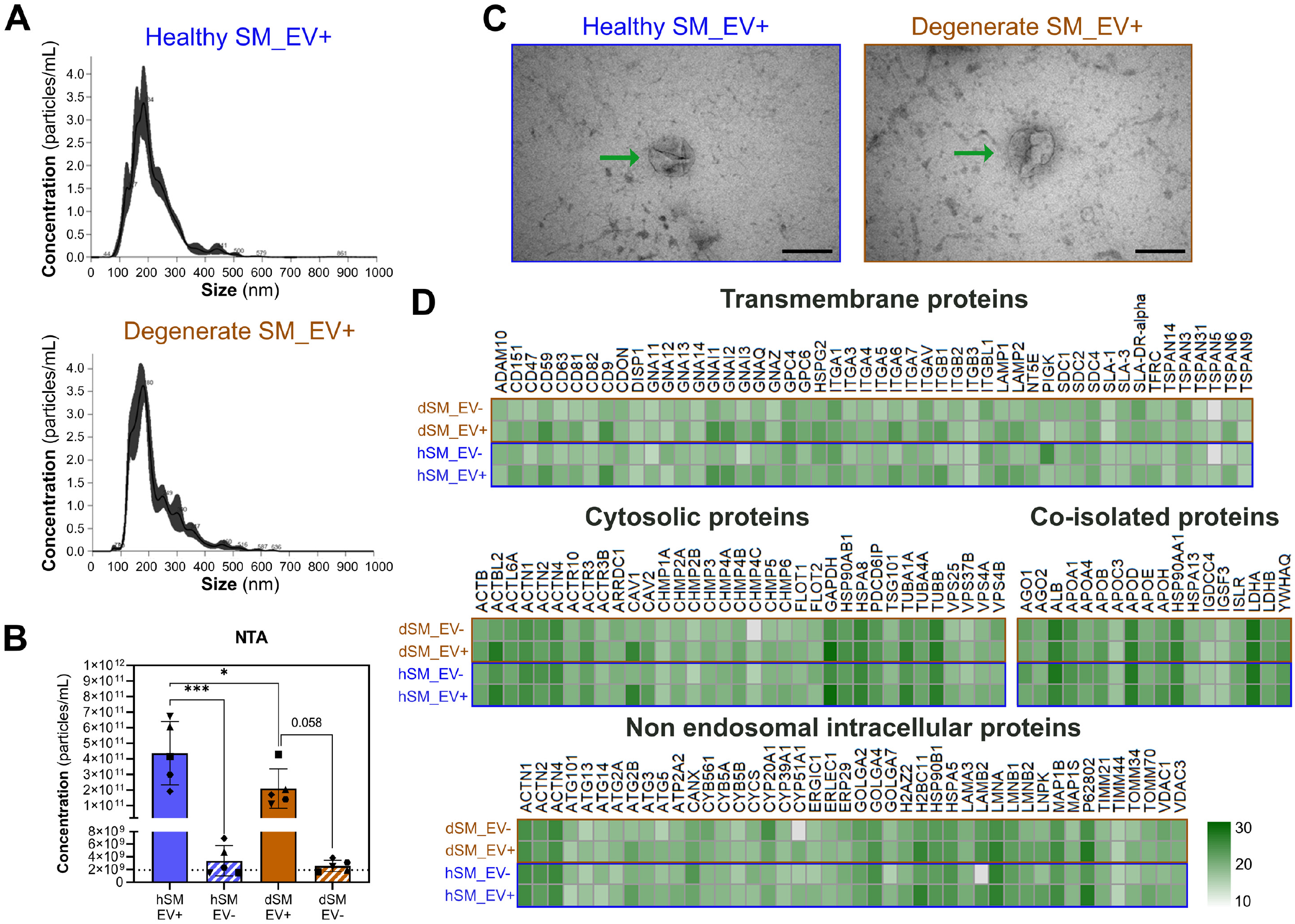
Characterization of notochordal cell extracellular vesicles (EVs) and their EV-depleted controls following culture in healthy or degenerate media. Characterization of EV-enriched secretome medium (SM_EV+) derived under a healthy (h) or degenerate (d) environment and their respective EV-depleted controls (SM_EV−). (**A**) Particle size distribution histograms of representative healthy (h)SM_EV+ and degenerate (d)SM_EV+. (**B**) Average particle concentration in hSM_EV+, hSM_EV−, dSM_EV+ and dSM_EV−. The dotted line represents plain DMEM controls as background. Dots indicate individual matched pig donor (N=5). Bars and whiskers indicate means ± SD. p-values are indicated for statistical significance of differences between conditions using Sidak’s multiple comparison test (numeric *p*-value: *p* < 0.1, * : *p* < 0.05, *** : *p* < 0.0005). (**C**) Transmission electron microscopy image of representative hSM_EV+ and dSM_EV− samples (scale bar represents 200 nm). (**D**) Normalized protein abundancy of classical EV markers in hSM_EV+, hSM_EV−, dSM_EV+ and dSM_EV−.

### GAGs are co-isolated with NC-EVs, with degenerative environment lowering the abundance of co-isolated IL10

Seeing the quantitative differences in NC-EVs released under healthy and degenerate conditions, and the fact that NCs reside within a matrix-rich tissue, we investigated whether the released GAGs differentially associated with NC-EVs by assessing the EV-enriched secretome media. hSM_EV+ and dSM_EV+ presented comparable GAG levels, however, they were around 10 times less rich in GAGs compared to the respective NCCM (**Figure 4C and 7D**). In the EV-depleted controls, GAG levels were significantly lower than those in SM_EV+ by 96% and 95%, respectively (**Figure 7D**). These results indicate that the GAGs released from the matrix-rich tissue into NCCM are largely removed upon the EV-enrichment procedure. The fraction of GAGs co-isolated with SM_EV+ was further reduced upon EV depletion, suggesting GAGs might be part of NC-EVs’ protein corona.

**Figure 7.**
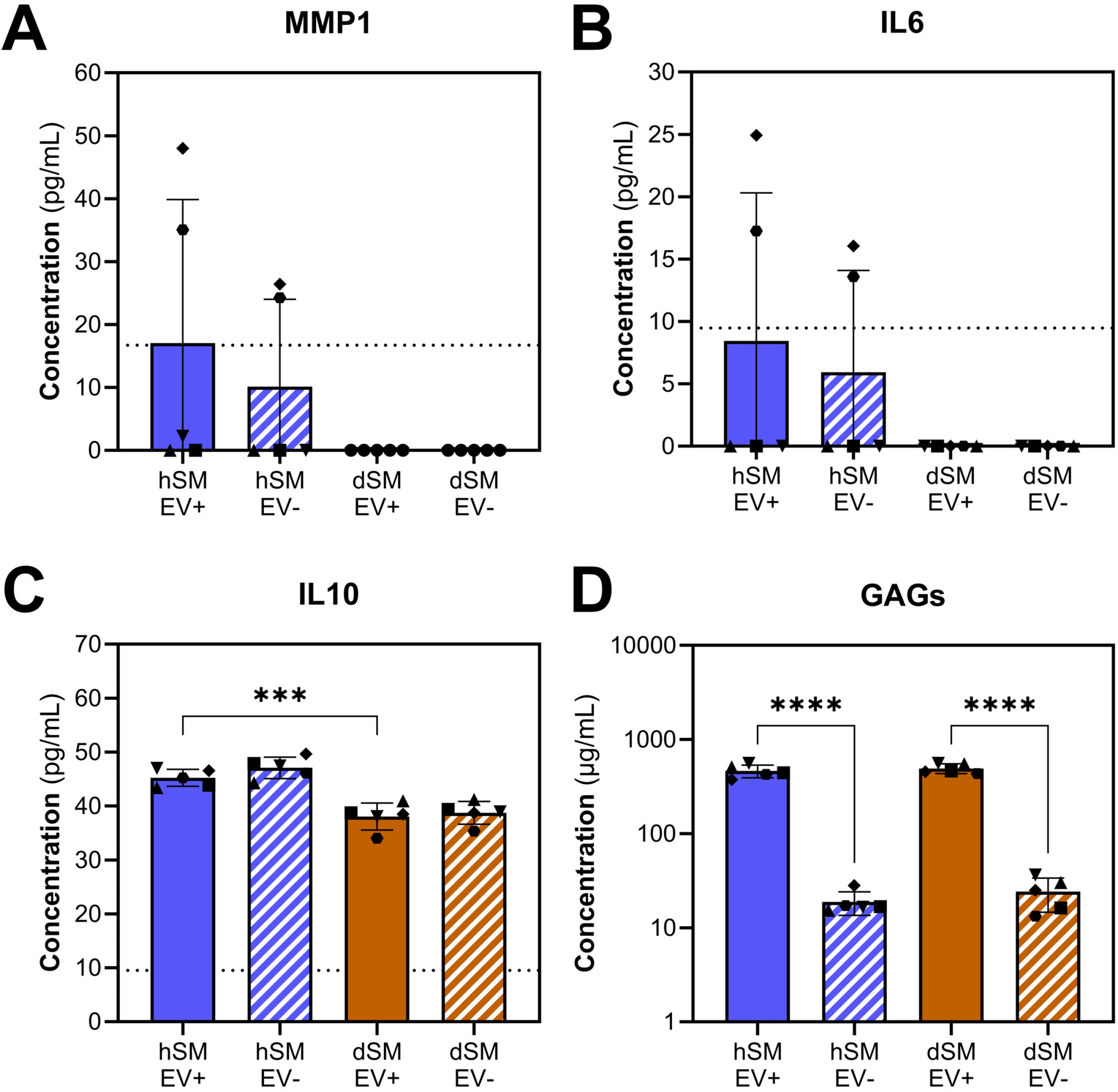
Co-isolated soluble factors in extracellular vesicle (EV) enriched secretome following culture in healthy or degenerate media. Measurements were done on EV-enriched secretome derived under healthy (hSM_EV+) or degenerate environment (dSM_EV+) and their respective EV-depleted controls (hSM_EV− and dSM_EV−). (**A-D**) Levels of MMP1, IL6, IL10, and GAGs in hSM_EV, hSM_EV−, dSM_EV+ and dSM_EV−. The dotted line represents the Multiplex ELISA detection limit of respective analytes (MMP1: 16.7 pg/mL; IL6: 9.5 pg/mL; IL10: 9.5 pg/mL). Dots indicate individual matched pig donor (N=5). Bars and whiskers indicate means ± SD. p-values are indicated for statistical significance of differences between conditions using Friedman test for MMP1 and IL6 graphs and Sidak’s multiple comparison test for IL10 and GAGs graphs (*** : *p* < 0.0005, **** : *p* < 0.0001).

Soluble proteins have been shown to bind to GAGs and contribute to the protein corona (Crijns et al., 2020; Lv et al., 2016). Therefore, we measured the levels of inflammatory regulators previously found in NCCM in SM_EV+ samples and interpreted the data in the context of the respective EV-depleted controls. Contrary to what was observed for NCCM samples, upon EV enrichment, MMP1 and IL6 levels became detectable in two independent hSM_EV+ samples and their respective hSM_EV− controls; however, they were still undetectable in all dSM_EV+ and SM_EV−samples (**Figure 7A-7B**). Consistent with trends observed for NCCM, but with a 10-fold enrichment in NC-EV samples, IL10 levels were lower in dSM_EV+ compared to hSM_EV+, and were comparable to their respective EV-depleted controls (**Figure 7C**). In addition, as with the previous findings on NCCM, TNF, IL1B, IL1RN, IFNG, and CXCL8 levels upon EV enrichment remained undetectable in all conditions (**Supplementary figure 1C-1D**).

Altogether, these results indicate that EV isolation from NCCM enriches MMP1, IL6 and IL10, while largely excludes soluble GAGs. EV-depletion also removed most of the soluble GAGs, while MMP1, IL6 and IL10 remained unaffected. These findings suggest that these inflammatory regulators are not tightly bound to the NC-EV-associated GAGs and could be part of the NC-EV soft corona.

### A degenerative environment impairs NC-secretome and NC-EVs modulation of NPC matrix and phenotype

To determine whether changes in the NC-EV enriched secretome (SM_EV+) translate into functional differences, we assessed whether they improved the degenerative phenotype of mildly degenerated NPCs and promoted anabolic regenerative responses (**Figure 1**). To mimic the degenerated disc environment, NPCs pellets were cultured in low glucose media (1 g/L) with pH 6.8 and 350 mOsm for 14 days (**Supplementary figure 2A**). EV-depleted controls (SM_EV−) were used to determine whether the observed effects of the secretome were EV-mediated.

To explore whether SM_EV+ modulates the cellular phenotype in an EV-mediated manner, we analysed NPC pellet characteristics and NPC marker expression following 14 days of SM_EV+ and SM_EV− treatment. No differences were observed in pellet size among treatments (**Figure 8A-9B**). However, DNA content in NPC pellets increased significantly with hSM_EV+ and tended to increase with dSM_EV+ (*p*=0.08), while it was comparable between their corresponding EV-depleted controls (**Figure 8C**). Surprisingly, overall TBXT (cytoplasmic and nuclear) immunopositivity in NPC pellets decreased with both hSM_EV+ and dSM_EV+ treatment. Reduced nuclear TBXT immunopositivity indicated reduced TBXT activity following both SM_EV+ treatments (**Figure 8D-9F**). Moreover, overall TBXT immunopositivity was only significantly lower in NPC pellets treated with hSM_EV+ than in hSM_EV− controls, and this was also confirmed for nuclear TBXT immunopositivity (*p*=0.1; **Figure 8D-F**). To expand on this observation, we expanded our analysis to additional cell markers. TIE2 levels in NPC pellets decreased following both SM_EV+ treatments and was comparable among conditions and EV-depleted controls (**Figure 8G** and **8H**), while KRT8/18/19 was maintained (**Figure 8I** and **9J**). These findings suggest that hNC-EVs mediated the reduction in TBXT abundance in NPCs, while the reduced TBXT levels in dSM_EV+ treated NPCs could not be attributed specifically to the dNC-EVs.

**Figure 8.**
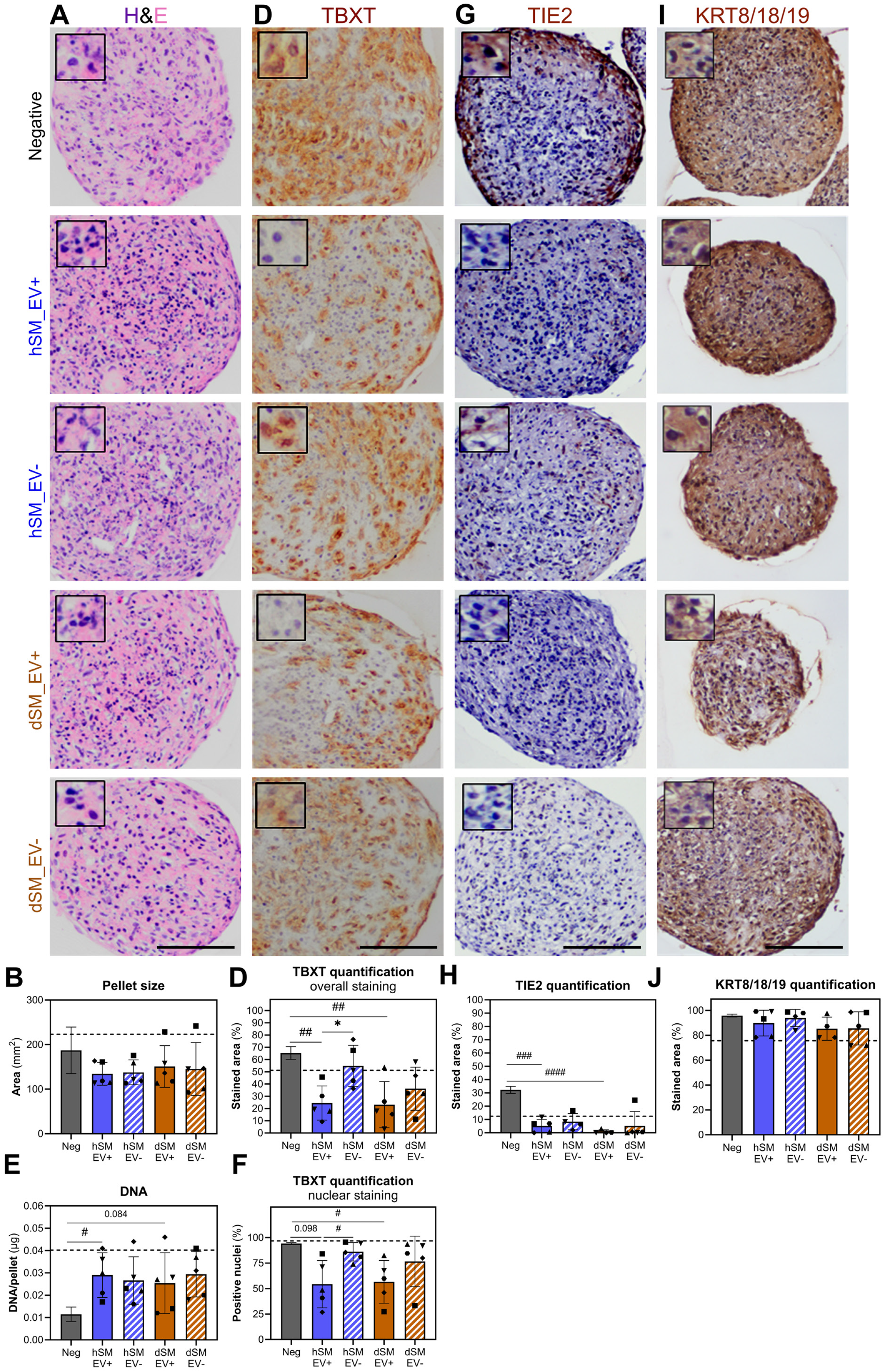
Functional impact of the extracellular vesicle (EV) enriched secretome derived from notochordal cells on nucleus pulposus cellularity and phenotype. All analysis were done on dog nucleus pulposus cell (NPC) pellets cultured for 14 days either in degenerate disc medium (Neg), or degenerate disc medium supplemented with hSM_EV+, hSM_EV−, dSM_EV+ or dSM_EV−. The NC-EV donor-specific effects were tested on a representative pooled dog NPC population (N=7). (**A**) Haematoxylin/Eosin (H&E) staining of NPC pellet (scale bar represents 100 µm). (**B-C**) Size and DNA content of NPC pellet upon 14 days treatment. (**D-J**) Immunohistochemistry for brachyury (TBXT), angiopoietin-1 receptor (TIE2) and cytokeratin 8+18+19 (KRT8+18+19) with corresponding quantification of percentage of positive stained area and of TBXT+ nuclei (scale bar represents 100 µm). Dots indicate individual matched pig donor-derived NC-EV treatment (N=5; n=3). Bars and whiskers indicate means ± SD. The dotted line represents levels in the positive (10 ng/mL TGF-β1) condition. *p*-values are indicated for statistical significance of differences among healthy and degenerate SM_EV+ and their respective SM_EV− conditions using Sidak’s multiple comparison test (numeric *p*-value: *p* < 0.1,* : *p* < 0.05) and between negative and hSM_EV+ and dSM_EV+ conditions using Welch’s unpaired t-test (numeric *p*-value: *p* < 0.1, # : *p* < 0.05, ## : *p* < 0.005, ### : *p* < 0.0005, #### : *p* < 0.0001).

**Figure 9.**
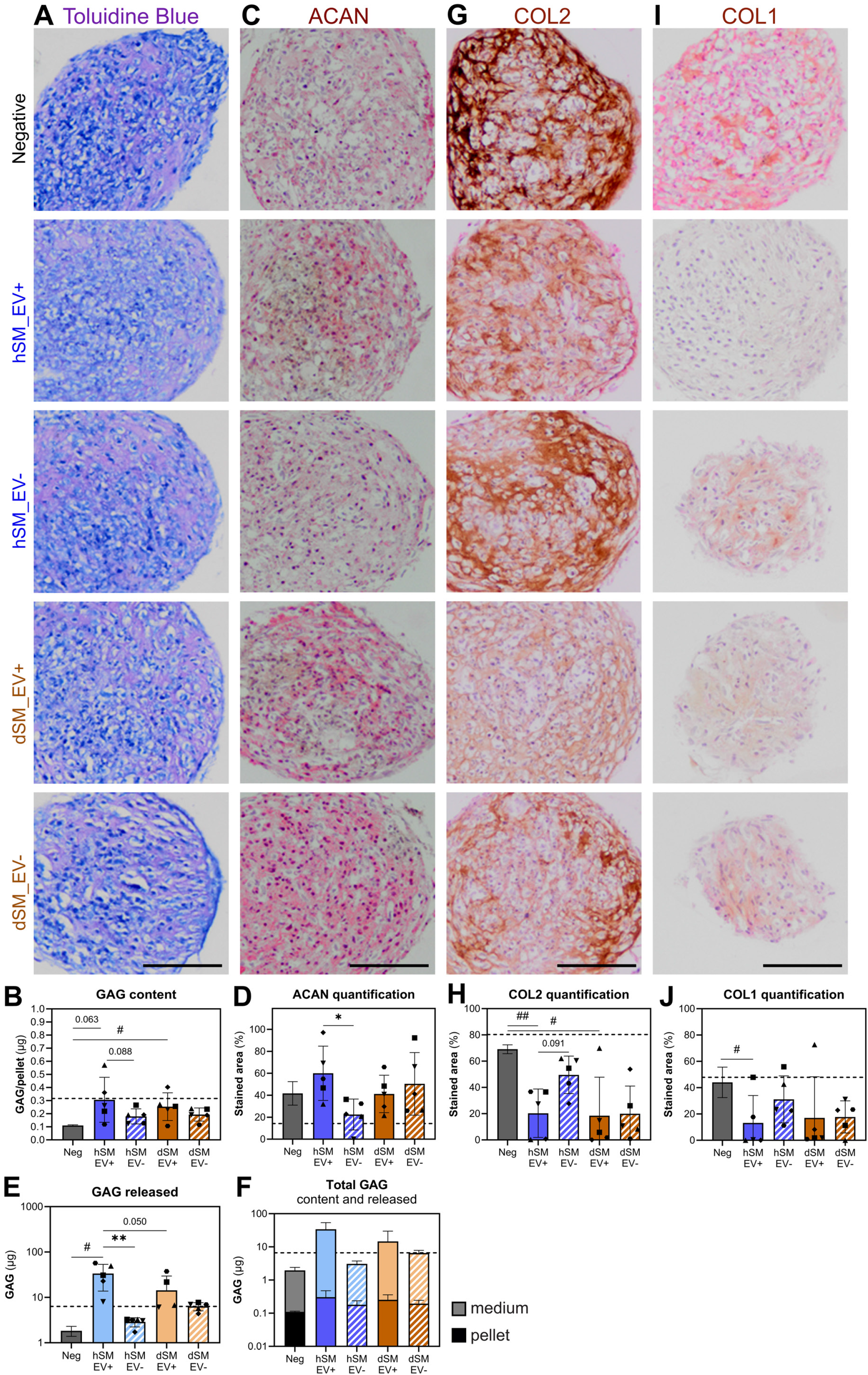
Functional impact of the extracellular vesicle (EV) enriched secretome derived from notochordal cells on nucleus pulposus cellularity and phenotype. All analyses were performed on dog nucleus pulposus (NPC) pellets cultured for 14 days in degenerate disc medium (Neg), degenerate disc medium supplemented with hSM_EV+, hSM_EV−, dSM_EV+ or dSM_EV−. The NC-EV donor-specific effects were tested on a representative pooled dog NPC population (N=7). (**A**) Toluidine Blue staining of NPC pellets (scale bar represents 100 µm). (**B**) Glycosaminoglycans (GAG) content of NPC pellet upon 14 days of treatment. (C-D) Immunohistochemistry for aggrecan (ACAN) with corresponding quantification of the percentage of positive stained area. (**E-F**) Cumulative GAG amount released from the pellet into the culture medium during 14 days of culture and respective total GAG amount = GAG content (pellet) + GAG released (medium). (**G-J**) Immunohistochemistry for collagen 2 (COL2) and collagen 1 (COL1) with corresponding quantification of percentage of positive area (scale bar represents 100 µm). Dots indicate individual matched pig donor-derived NC-EV treatment (N=5; n=3). Bars and whiskers indicate mean ± SD. The dotted line represents levels in the positive (10 ng/mL TGF-β1) condition. *p*-values are indicated for statistical significance among hSM_EV+ and dSM_EV+ and their respective hSM_EV− and dSM_EV− conditions using Sidak’s multiple comparison test (numeric *p*-value: *p* < 0.1, * : *p* < 0.05, ** : *p* < 0.005) and between negative and hSM_EV+ or dSM_EV+ conditions using Welch’s unpaired t-test (numeric *p*-value: *p* < 0.1, # : *p* < 0.05, ## : *p* < 0.005).

Next, we assessed whether these phenotypic changes are reflected in distinct ECM profiles. The GAG content of NPC pellets increased comparably following stimulation with hSM_EV+ (*p*=0.06) and dSM_EV+ (*p*=0.04) compared to the negative control (**Figure 9A** and **9B**). NPC pellets treated with hSM_EV+ tended to have a higher GAG content compared to SM_EV− controls (*p*=0.088), while this was not the case for NPC pellets treated with dSM_EV+ compared to dSM_EV− controls (**Figure 9A** and **9B**). This difference was further corroborated by higher immunopositivity for ACAN (**Fig 9C** and **9D**), indicating hNC-EVs augmented ACAN deposition in the NPC pellets. To calculate the total GAG production during culture, we analysed the GAGs released in the media. GAG release increased following treatment with SM_EV+. This increase was significant in hSM_EV+ (*p*=0.02) group, but did not reach statistical significance (*p*=0.14) in dSM_EV+ (**Figure 9E**). Cumulatively, NPC pellets treated with hSM_EV+ tended (*p*=0.05) to release 2.4-fold more GAGs compared to NPC pellets treated with dSM_EV+ (**Figure 8E**). GAG levels released and the estimated total GAG production (**Figure 9F**) by NPC pellets were higher following hSM_EV+ vs. SM_EV− treatment, while there was no difference between dSM_EV+ and dSM_EV− treatment (**Figure 9E**). To investigate whether SM_EV+ influenced other ECM components, we assessed COL2 and COL1 immunopositivity. COL2 and COL1 levels in NPC pellets were significantly decreased following hSM_EV+ and dSM_EV+ treatment compared to the negative control, with the decrease in COL1 levels following dSM_EV+ failing to reach statistical significance (*p*=0.13; **Figure 9G-8J**). Moreover, while COL2 and COL1 levels in NPC pellets were largely comparable between SM_EV+ and corresponding SM_EV− stimulation (**Figure 9G-8J**), a trend towards lower COL2 levels was observed following hSM_EV+ treatment compared with SM_EV− (*p*=0.09; **Figure 9I** and **9J**).

Overall, these results indicate that hNC-EVs enriched in hSM_EV+ media improved GAG production and deposition and reduced COL2 deposition by NPCs, while this EV-mediated effect appeared to diminish after treatment with EV-enriched secretome generated under degenerate conditions. We confirmed the role of NC-EVs in matrix deposition by serial dilution studies of SM_EV+ (**Supplementary figure 4**).

## Discussion

Changes in the nutrient-limited IVD environment, progressively shifting towards acidic and hypo-osmolar conditions, are considered among the key environmental factors contributing to the pathophysiological processes of IVD degeneration (Bibby et al., 2005). Alongside this, at the cellular level, the large vacuolated NC population transitions to the smaller non-vacuolated NPCs (Bach et al., 2022). The present study assessed whether this degenerative environment influences the NC secretome and modulates NC-EV release. Using mild degenerate NPCs, we studied the functional role of the NC-secretome in regulating ECM production and cell phenotype maintenance *in vitro*, and assessed NC-EV-mediated effects by comparing SM_EV+ to SM_EV− methodological controls. We show that the degenerative environment reduced EV release by NCs, while maintaining similar levels of inflammatory mediators associated with the soft corona, except for lower IL-10 levels in degenerative conditions. Overall, the NC-secretome supported pellet viability, as evidenced by increased DNA levels across all treatment conditions, and with NPCs shifting from a progenitor state to more mature matrix producing cells expressing lower levels of TBXT and TIE2. Although both healthy and degenerate SM_EV+ elicited biological activity, only hSM_EV+ demonstrated EV-specific activity as evidenced by an increased GAG release and ACAN deposition and reduced TBXT levels and COL2 deposition.

The observed negating effects of the degenerative environment on NC-EV-mediated phenotypic transition, together with the fact that EVs closely resemble the status of their source (Almeria et al., 2022), urged us to study the effect of the degenerative environment on the NC-rich NP tissue. The short culture period in low-glucose media for 4 days led to subtle changes in the NC phenotype, while maintaining tissue viability and common NC markers, TBXT and KRT8/18/19 protein levels were comparable to native tissue levels. NCs are notorious for losing their cell phenotype (Williams et al., 2023), e.g. even immediately after cell isolation, and only partly recover *TBXT* and *KRT18* gene expression over 28 days when cultured in alginate (Arkesteijn et al., 2015), with low glucose and adjusted osmolarity benefiting *TBXT* and *KRT8/18* expression in dog NC clusters (Spillekom et al., 2014; Arkesteijn et al., 2015). This seems to contradict our finding where TBXT and KRT8/18/19 expression was comparable between low glucose hypo- and hyperosmotic conditions, but it may be due to the much shorter tissue culture period. Overall, NC markers were shown to be largely preserved during short NC tissue culture in low glucose, either healthy or degenerate conditions.

Following short-term tissue culture, in both culture environments the size of the NC vacuole, which commonly occupy most of the cell volume, increased compared to the native NC-tissue, with larger vacuoles under healthy compared to degenerative conditions. This is in contrast with previous findings, showing that NCs maintain their cell volume during hypotonic stress via their vacuoles, which have an osmoregulatory function and release their hypo-osmotic content to maintain osmotic homeostasis (Hunter et al., 2007). We therefore attribute the increased cell volume in the present study to the large swelling capacity of the GAG-rich NP tissue under free swelling conditions (Salzer et al., 2022). Similarly, in bovine tissues, free swelling leads to matrix remodelling and GAG release into the culture media, resulting in a local pericellular hypotonic microenvironment causing the cells to swell (Mizuno et al., 2024). In line with this, in the present study ∼80% of the GAGs was released into the media, both under healthy and degenerative conditions, resulting in reduced ACAN deposition and decreased toluidine blue metachromasia. Despite these changes, deposited COL2 was maintained and COL1 remained undetectable in both healthy and degenerate conditions, probably because of the nature of collagen fibers that form a more stable network than GAGs (reviewed by Kiani et al., 2002; Wei et al., 2019). These results are consistent with previous research showing that NP tissue cultured for 7 days in high glucose, either under hypo or hyperosmolar conditions, maintained COL2 expression, while COL1 expression remained absent (Li et al., 2016; Mizuno et al., 2024). Altogether, these findings indicate that short culture of NC tissue decreased the GAG-rich matrix, contributing to a decrease in the surrounding osmotic pressure causing an increase in the intra-cellular NC vacuoles size.

The present study investigated whether the degenerative environment influenced NC biology and functional response to low pH and osmolarity. Following short NC-tissue culture, degenerative stimuli did not increase the release of the investigated pro-inflammatory mediators, except for the lower PGE2 release compared to healthy culture conditions. Similarly, human degenerate NP tissues cultured in healthy or degenerate media exhibited only mild modulations in a subset of inflammatory mediators (i.e.. CCL4, CXCL10) amongst a multiplex panel after the addition of 100 pg/mL IL-1β to the degenerate media (Snuggs et al., 2025). PGE2 has been primarily studied as a catabolic and inflammatory mediator (Vo et al., 2010). To date, the effect of low pH on PGE2 has not been reported. Studies with bovine NPC-rich NP tissue showed that hypo-osmotic culture conditions result in lower PGE2 release compared to hyperosmotic culture conditions (Mouser et al., 2019), while dynamic compressive and hyper-physiological loading of bovine NP cells embedded in an agarose-collagen hydrogel led to increased PGE2 release (Cambria et al., 2021). Overall, these findings indicate that the combination of low pH and osmolarity minimally affected NCs which are maintained in their native tissue and seemed only to temper PGE2 release.

Our findings show that a degenerative environment defined by low pH, combined with hypo-osmolar media, and a changing tissue landscape in which the pericellular matrix becomes gradually GAG-depleted, reduced the number of NC-EVs released, without affecting their size and morphology. Previous studies on other species and cell types using monolayer cell culture, such as human fibroblasts, highlight that osmotic stress promotes EV release (Parvanian et al., 2021). Moreover, pH 5 in HeLa cell culture decreased EV release by impairing the intracellular membrane mobility required for EV production and release (Nakase et al., 2021). The reduced NC-EV release is unlikely due to differences in retention of the EVs within the ECM, as both healthy and degenerative culture conditions showed comparable matrix composition and degradation of the GAG-rich tissue over time. Inflammatory mediators co-isolated with the EV fractions were likewise largely comparable between the two conditions, with notably lower levels of the anti-inflammatory mediator IL-10 under degenerative culture conditions. Altogether, our study indicates that a degenerative environment reduces the release of NC-EVs and specific co-isolates, suggesting that the degenerate NC-EVs might present an altered biomolecular corona composition.

To better resemble the nutrient-deprived IVD environment, we used low-glucose media, and an adjusted pH and osmolarity, to generate SM_EV+ and to study their biological functionality on NPC pellets, instead of high-glucose conditions used previously (Bach et al., 2017; Bach et al., 2016; van Maanen et al., 2025). This lead to findings that were partly consistent with, and partly opposite, to our earlier results. The SM_EV+ increased DNA content, indicative of improved viability, and matrix NPC pellet content, while not affecting ACAN expression, albeit reaching lower absolute levels compared to NPC pellets cultured in high-glucose conditions and treated with SM_EV+ generated in high glucose media not adjusted for pH or osmolarity (Bach et al., 2017; van Maanen et al., 2025). This difference could be attributed to the lower metabolic activity of NPCs under low-glucose conditions (Bibby et al., 2005; Huang et al., 2007). Furthermore, the SM_EV+ reduced the expression of the healthy matrix component COL2 and the fibrotic marker COL1 compared to untreated controls. These findings contrast previous reports showing that SM_EV+ derived under high-glucose conditions did not affect *ACAN*, *COL2A1* or *COL1A1* gene expression (van Maanen et al., 2025). This difference might be explained by the different glucose concentrations of the two experimental setups, where in the present study lower glucose levels are presumably responsible for changes in the NC-secretome composition, as well as for a different response of NPC to such stimuli, towards cell survival over energy-consuming matrix remodelling (Guehring et al., 2009; Naqvi & Buckley, 2015; Rinkler et al., 2010). Interestingly, healthy and degenerate SM_EV+ produced comparable increases in NPC pellet DNA and matrix content, as well as comparable inhibitory effects on COL2 and COL1 expression, while healthy SM_EV+ induced higher GAG release compared to the degenerate SM_EV+. This finding is the net balance of matrix production, release and degradation, leading to an increase in matrix release. Which of these processes is leading remains to be determined. Moreover, we dissected EV-mediated effects from SM_EV+ using an EV-depleted methodological control and found that the increased GAG release by NPC pellets, as well as decreased COL2 expression, were shown to be partly EV-mediated only for the healthy NC-secretome. There are several caveats to consider. The study was designed to investigate the environmental effects on the EV-enriched NC-secretome, and therefore performed functional studies using SV_EV+ normalised for starting tissue quantities when generating conditioned media. Taking this into consideration, the absence of a detectable De-EV-mediated effect may be attributable to several factors. First, the degenerative environment attenuated EV release, thereby limiting the biological activity of EV-mediated effects of SM_EV+. Second, because the EVs were isolated from a complex conditioned medium, bioactive non-EV secretome components may have masked EV-specific effects. Finally, the employed measures may not have been sufficiently sensitive to detect subtle EV-mediated responses. Taken together, these findings suggest that low pH and osmolarity negate the beneficial matrix anabolic effects of the NC-secretome, and that the observed effects are only partly EV-mediated.

This negating influence of the degenerative IVD environment might be explained by the reduced EV release, and/or changes in the EV cargo composition, and therefore biological function. The EV cargo of human NP cells derived from degenerated disc tissue has previously been shown to change, becoming enriched with proteins involved in vesicle-mediated transport and matrix structure and organization (Li et al., 2024). In this perspective, the observed lower IL10 levels co-isolated with dSM_EV+ in the present study could contribute to the lower matrix anabolic capacity. IL10 plays a positive role in maintaining ECM homeostasis and alleviates IVD degeneration (Ge et al., 2020; Holm et al., 2009). Notably, the majority of analysed inflammation molecules was not detected in association with SM_EV+ upon EV enrichment, which implies that these did not contribute to EV-mediated effects. Also, the levels of GAG co-isolated with SM_EV+ were drastically lower upon EV-depletion, indicating that GAGs associated to NC-EVs and contributed to their matrix-rich protein corona. The presence of GAGs in SM_EV+ partly explains the observed anabolic effects observed on NPC pellets (de Vries et al., 2019; Illien-Jünger et al., 2016). Further studies are needed to understand how low pH and osmolarity influence the composition and function of SM_EV+ and the protein corona.

The diminished effects of SM_EV+ at the matrix level were corroborated by the NPC phenotype. To the best of our knowledge, effects of SM_EV+ on the NPC phenotype have not been previously described. Both healthy and degenerate SM_EV+ decreased the expression of TBXT and TIE2, while preserving KRT8/18/19, implying a phenotypic maturation shift towards matrix-producing NPCs. Notably, the reduction of TBXT expression in NPCs was only EV-mediated in hSM_EV+. The observed effects cannot be clarified exclusively to TBXT or TIE2-dependent signalling, since anabolic pathways such as TGF-β signaling may also apply (Chen et al., 2019). Elucidating the underlying mechanisms require a more comprehensive characterization of the culture medium to identify the bioactive factors responsible for these effects.

In conclusion, this study shows that early disc degenerative processes, in an ECM becoming GAG-depleted, affect the NC secretome, contributing to the NC-to-NPC transition; and decreases in pH and osmolarity attenuate the release of NC-EVs. They are also shown to impair the EV-mediated capacity of the NC secretome to promote healthy matrix production, negating the beneficial role of NC-EVs in IVD health and homeostasis. It remains to be determined whether the observed effects are solely attributable to changes in NC-EV abundance or whether alterations in NC-EV cargo and the associated biomolecular corona occur in response to changes in ECM composition and the microenvironment.

## Acknowledgements

We thank Cornelis Seinen for TEM microscopy, Paula Sobrevals Alcaraz for LC/MS data processing, dr. Michelle Teunissen for technical assistance with multiplex ELISA. We are thankful to the UMC Utrecht Proteomics facility, which is part of the Oncode Accelerator Project that has received funding from the Dutch National Growth Fund (NGF) under grant number NGFOP2201, for support with the mass spectrometry. Figures 1 and 2 were prepared using BioRender.

## Contributions

Conceptualization: D.C., C.V., M.H.M.W., M.A.T.; Methodology: D.C., C.V., P.V. H.R.V.; Investigation: D.C., C.V., A.E.; Data curation: D.C., C.V., F.M.R.; Formal analysis: D.C., C.V., M.H.M.W., M.A.T.; Resources: P.V., H.R.V., M.H.M.W., M.A.T.; Writing original draft: D.C., C.V., M.A.T.; Writing, review and editing: all authors; Supervision: C.V., K.I., M.H.M.W., M.A.T.; Funding acquisition: M.A.T.

## Declaration of Generative AI in scientific writing

During the preparation of this manuscript the authors used Grammarly and ChatGPT (version GPT-5.5, OpenAI) for assistance with grammar, spelling and readability of the manuscript. After using these tools the authors reviewed and edited the content as needed and take full responsibility for the content.

## References

Adams, M. A., & Roughley, P. J. (2006). What is intervertebral disc degeneration, and what causes it? Spine, 31(18), 2151–2161. 10.1097/01.brs.0000231761.73859.2c

Almeria, C., Kreß, S., Weber, V., Egger, D., & Kasper, C. (2022). Heterogeneity of mesenchymal stem cell-derived extracellular vesicles is highly impacted by the tissue/cell source and culture conditions. Cell & Bioscience, 12(1), 51. 10.1186/s13578-022-00786-7

Arkesteijn, I. T. M., Smolders, L. A., Spillekom, S., Riemers, F. M., Potier, E., Meij, B. P., Ito, K., & Tryfonidou, M. A. (2015). Effect of coculturing canine notochordal, nucleus pulposus and mesenchymal stromal cells for intervertebral disc regeneration. Arthritis Research & Therapy, 17(1), 60. 10.1186/s13075-015-0569-6

Bach, de Vries, S. A., Riemers, F. M., Boere, J., van Heel, F. W., van Doeselaar, M., Goerdaya, S. S., Nikkels, P. G., Benz, K., Creemers, L. B., Maarten Altelaar, A. F., Meij, B. P., Ito, K., & Tryfonidou, M. A. (2016). Soluble and pelletable factors in porcine, canine and human notochordal cell-conditioned medium: Implications for IVD regeneration. European Cells & Materials, 32, 163–180. 10.22203/ecm.v032a11

Bach, F., Libregts, S., Creemers, L., Meij, B., Ito, K., Wauben, M., & Tryfonidou, M. (2017). Notochordal-cell derived extracellular vesicles exert regenerative effects on canine and human nucleus pulposus cells. Oncotarget, 8(51), 88845–88856. 10.18632/oncotarget.21483

Bach, Poramba-Liyanage, D. W., Riemers, F. M., Guicheux, J., Camus, A., Iatridis, J. C., Chan, D., Ito, K., Le Maitre, C. L., & Tryfonidou, M. A. (2022). Notochordal Cell-Based Treatment Strategies and Their Potential in Intervertebral Disc Regeneration. Frontiers in Cell and Developmental Biology, 9. 10.3389/fcell.2021.780749

Bartels, E. M., Fairbank, J. C., Winlove, C. P., & Urban, J. P. (1998). Oxygen and lactate concentrations measured in vivo in the intervertebral discs of patients with scoliosis and back pain. Spine, 23(1), 1–7; discussion 8. 10.1097/00007632-199801010-00001

Bibby, S. R. S., Jones, D. A., Ripley, R. M., & Urban, J. P. G. (2005). Metabolism of the Intervertebral Disc: Effects of Low Levels of Oxygen, Glucose, and pH on Rates of Energy Metabolism of Bovine Nucleus Pulposus Cells. Spine, 30(5), 487. 10.1097/01.brs.0000154619.38122.47

Chen, S., Liu, S., Ma, K., Zhao, L., Lin, H., & Shao, Z. (2019). TGF-β signaling in intervertebral disc health and disease. Osteoarthritis and Cartilage, 27(8), 1109–1117. 10.1016/j.joca.2019.05.005

Cheung, K. M. C., Karppinen, J., Chan, D., Ho, D. W. H., Song, Y.-Q., Sham, P., Cheah, K. S. E., Leong, J. C. Y., & Luk, K. D. K. (2009). Prevalence and Pattern of Lumbar Magnetic Resonance Imaging Changes in a Population Study of One Thousand Forty-Three Individuals. Spine, 34(9), 934. 10.1097/BRS.0b013e3181a01b3f

Crijns, H., Vanheule, V., & Proost, P. (2020). Targeting Chemokine—Glycosaminoglycan Interactions to Inhibit Inflammation. Frontiers in Immunology, 11, 483. 10.3389/fimmu.2020.00483

de Vries, S., Doeselaar, M. van, Meij, B., Tryfonidou, M., & Ito, K. (2019). Notochordal Cell Matrix As a Therapeutic Agent for Intervertebral Disc Regeneration. Tissue Engineering Part A, 25(11–12), 830–841. 10.1089/ten.tea.2018.0026

Farndale, R. W., Sayers, Christine A., & and Barrett, A. J. (1982). A Direct Spectrophotometric Microassay for Sulfated Glycosaminoglycans in Cartilage Cultures. Connective Tissue Research, 9(4), 247–248. 10.3109/03008208209160269

Ferreira, M. L., de Luca, K., Haile, L. M., Steinmetz, J. D., Culbreth, G. T., Cross, M., Kopec, J. A., Ferreira, P. H., Blyth, F. M., Buchbinder, R., Hartvigsen, J., Wu, A.-M., Safiri, S., Woolf, A. D., Collins, G. S., Ong, K. L., Vollset, S. E., Smith, A. E., Cruz, J. A., … March, L. M. (2023). Global, regional, and national burden of low back pain, 1990–2020, its attributable risk factors, and projections to 2050: A systematic analysis of the Global Burden of Disease Study 2021. The Lancet Rheumatology, 5(6), e316–e329. 10.1016/S2665-9913(23)00098-X

Ge, J., Yan, Q., Wang, Y., Cheng, X., Song, D., Wu, C., Yu, H., Yang, H., & Zou, J. (2020). IL-10 delays the degeneration of intervertebral discs by suppressing the p38 MAPK signaling pathway. Free Radical Biology & Medicine, 147, 262–270. 10.1016/j.freeradbiomed.2019.12.040

Guehring, T., Wilde, G., Sumner, M., Grünhagen, T., Karney, G. B., Tirlapur, U. K., & Urban, J. P. G. (2009). Notochordal intervertebral disc cells: Sensitivity to nutrient deprivation. Arthritis & Rheumatism, 60(4), 1026–1034. 10.1002/art.24407

Guerrero, J., Häckel, S., Croft, A. S., Albers, C. E., & Gantenbein, B. (2021). The effects of 3D culture on the expansion and maintenance of nucleus pulposus progenitor cell multipotency. JOR Spine, 4(1), e1131. 10.1002/jsp2.1131

Hodson, N. W., Patel, S., Richardson, S. M., Hoyland, J. A., & Gilbert, H. T. J. (2018). Degenerate intervertebral disc-like pH induces a catabolic mechanoresponse in human nucleus pulposus cells. JOR Spine, 1(1), e1004. 10.1002/jsp2.1004

Holm, S., Mackiewicz, Z., Holm, A. K., Konttinen, Y. T., Kouri, V.-P., Indahl, A., & Salo, J. (2009). Pro-inflammatory, pleiotropic, and anti-inflammatory TNF-α, IL-6, and IL-10 in experimental porcine intervertebral disk degeneration. Veterinary Pathology, 46(6), 1292–1300. 10.1354/vp.07-VP-0179-K-FL

Huang, C.-Y. C., Yuan, T.-Y., Jackson, A. R., Hazbun, L., Fraker, C., & Gu, W. Y. (2007). Effects of low glucose concentrations on oxygen consumption rates of intervertebral disc cells. Spine, 32(19), 2063–2069. 10.1097/BRS.0b013e318145a521

Hughes, C. S., Foehr, S., Garfield, D. A., Furlong, E. E., Steinmetz, L. M., & Krijgsveld, J. (2014). Ultrasensitive proteome analysis using paramagnetic bead technology. Molecular Systems Biology, 10(10), MSB145625. 10.15252/msb.20145625

Hunter, C. J., Bianchi, S., Cheng, P., & Muldrew, K. (2007). Osmoregulatory function of large vacuoles found in notochordal cells of the intervertebral disc running title: An osmoregulatory vacuole. Molecular & Cellular Biomechanics: MCB, 4(4), 227–237.

Illien-Jünger, S., Sedaghatpour, D. D., Laudier, D. M., Hecht, A. C., Qureshi, S. A., & Iatridis, J. C. (2016). Development of a Bovine Decellularized Extracellular Matrix-Biomaterial for Nucleus Pulposus Regeneration. Journal of Orthopaedic Research : Official Publication of the Orthopaedic Research Society, 34(5), 876–888. 10.1002/jor.23088

Ishihara, H., Warensjo, K., Roberts, S., & Urban, J. P. (1997). Proteoglycan synthesis in the intervertebral disk nucleus: The role of extracellular osmolality. American Journal of Physiology-Cell Physiology, 272(5), C1499–C1506. 10.1152/ajpcell.1997.272.5.C1499

Jackson, A., Huang, C.-Y., Brown, M., & Gu, W. (2011). 3D Finite Element Analysis of Nutrient Distributions and Cell Viability in the Intervertebral Disc: Effects of Deformation and Degeneration. Journal of Biomechanical Engineering, 133, 091006. 10.1115/1.4004944

Kiani, C., Chen, L., Wu, Y. J., Yee, A. J., & Yang, B. B. (2002). Structure and function of aggrecan. Cell Research, 12(1), 19–32. 10.1038/sj.cr.7290106

Laagland, L. T., Bach, F. C., Creemers, L. B., Le Maitre, C. L., Poramba-Liyanage, D. W., & Tryfonidou, M. A. (2022). Hyperosmolar expansion medium improves nucleus pulposus cell phenotype. JOR Spine, 5(3), e1219. 10.1002/jsp2.1219

Lan, W.-R., Pan, S., Li, H.-Y., Sun, C., Chang, X., Lu, K., Jiang, C.-Q., Zuo, R., Zhou, Y., & Li, C.-Q. (2019). Inhibition of the Notch1 Pathway Promotes the Effects of Nucleus Pulposus Cell-Derived Exosomes on the Differentiation of Mesenchymal Stem Cells into Nucleus Pulposus-Like Cells in Rats. Stem Cells International, 2019, 8404168. 10.1155/2019/8404168

Li, L., Al-Jallad, H., Sun, A., Georgiopoulos, M., Bokhari, R., Ouellet, J., Jarzem, P., Cherif, H., & Haglund, L. (2024). The proteomic landscape of extracellular vesicles derived from human intervertebral disc cells. JOR SPINE, 7(4), e70007. 10.1002/jsp2.70007

Li, P., Gan, Y., Xu, Y., Li, S., Song, L., Li, S., Li, H., & Zhou, Q. (2016). Osmolarity affects matrix synthesis in the nucleus pulposus associated with the involvement of MAPK pathways: A study of ex vivo disc organ culture system. Journal of Orthopaedic Research, 34(6), 1092–1100. 10.1002/jor.23106

Luoma, K., Riihimäki, H., Luukkonen, R., Raininko, R., Viikari-Juntura, E., & Lamminen, A. (2000). Low back pain in relation to lumbar disc degeneration. Spine, 25(4), 487–492. 10.1097/00007632-200002150-00016

Lv, Q., Zeng, J., & He, L. (2016). The advancements of heparanase in fibrosis. International Journal of Molecular Epidemiology and Genetics, 7(4), 137–140.

Matta, A., Karim, M. Z., Isenman, D. E., & Erwin, W. M. (2017). Molecular Therapy for Degenerative Disc Disease: Clues from Secretome Analysis of the Notochordal Cell-Rich Nucleus Pulposus. Scientific Reports, 7, 45623. 10.1038/srep45623

Mizuno, H. L., Kang, J. D., & Mizuno, S. (2024). Effects of hydrostatic pressure, osmotic pressure, and confinement on extracellular matrix associated responses in the nucleus pulposus cells *ex vivo*. Matrix Biology, 134, 162–174. 10.1016/j.matbio.2024.10.005

Mouser, V. H. M., Arkesteijn, I. T. M., van Dijk, B. G. M., Wuertz-Kozak, K., & Ito, K. (2019). Hypotonicity differentially affects inflammatory marker production by nucleus pulposus tissue in simulated disc degeneration versus herniation. Journal of Orthopaedic Research, 37(5), 1110–1116. 10.1002/jor.24268

Nakase, I., Ueno, N., Matsuzawa, M., Noguchi, K., Hirano, M., Omura, M., Takenaka, T., Sugiyama, A., Bailey Kobayashi, N., Hashimoto, T., Takatani-Nakase, T., Yuba, E., Fujii, I., Futaki, S., & Yoshida, T. (2021). Environmental pH stress influences cellular secretion and uptake of extracellular vesicles. FEBS Open Bio, 11(3), 753–767. 10.1002/2211-5463.13107

Naqvi, S. M., & Buckley, C. T. (2015). Extracellular matrix production by nucleus pulposus and bone marrow stem cells in response to altered oxygen and glucose microenvironments. Journal of Anatomy, 227(6), 757–766. 10.1111/joa.12305

O’Sullivan, K., O’Sullivan, P. B., & O’Keeffe, M. (2019). The Lancet series on low back pain: Reflections and clinical implications. British Journal of Sports Medicine, 53(7), 392–393. 10.1136/bjsports-2018-099671

Parvanian, S., Zha, H., Su, D., Xi, L., Jiu, Y., Chen, H., Eriksson, J. E., & Cheng, F. (2021). Exosomal Vimentin from Adipocyte Progenitors Protects Fibroblasts against Osmotic Stress and Inhibits Apoptosis to Enhance Wound Healing. International Journal of Molecular Sciences, 22(9), 4678. 10.3390/ijms22094678

Razaq, S., Wilkins, R. J., & Urban, J. P. G. (2003). The effect of extracellular pH on matrix turnover by cells of the bovine nucleus pulposus. European Spine Journal: Official Publication of the European Spine Society, the European Spinal Deformity Society, and the European Section of the Cervical Spine Research Society, 12(4), 341–349. 10.1007/s00586-003-0582-3

Ren, P., Chen, P., Reeves, R. A., Buchweitz, N., Niu, H., Gong, H., Mercuri, J., Reitman, C. A., Yao, H., & Wu, Y. (2023). Diffusivity of Human Cartilage Endplates in Healthy and Degenerated Intervertebral Disks. Journal of Biomechanical Engineering, 145(7), 071006. 10.1115/1.4056871

Richardson, S. M., Ludwinski, F. E., Gnanalingham, K. K., Atkinson, R. A., Freemont, A. J., & Hoyland, J. A. (2017). Notochordal and nucleus pulposus marker expression is maintained by sub-populations of adult human nucleus pulposus cells through aging and degeneration. Scientific Reports, 7(1), 1501. 10.1038/s41598-017-01567-w

Rinkler, C., Heuer, F., Pedro, M. T., Mauer, U. M., Ignatius, A., & Neidlinger-Wilke, C. (2010). Influence of low glucose supply on the regulation of gene expression by nucleus pulposus cells and their responsiveness to mechanical loading: Laboratory investigation. Journal of Neurosurgery: Spine, 13(4), 535–542. 10.3171/2010.4.SPINE09713

Risbud, M. V., Schoepflin, Z. R., Mwale, F., Kandel, R. A., Grad, S., Iatridis, J. C., Sakai, D., & Hoyland, J. A. (2015). Defining the phenotype of young healthy nucleus pulposus cells: Recommendations of the Spine Research Interest Group at the 2014 annual ORS meeting. Journal of Orthopaedic Research, 33(3), 283–293. 10.1002/jor.22789

Sadowska, A., Kameda, T., Krupkova, O., & Wuertz-Kozak, K. (2018). Osmosensing, osmosignalling and inflammation: How intervertebral disc cells respond to altered osmolarity. European Cells & Materials, 36, 231–250. 10.22203/eCM.v036a17

Sakai, D., Schol, J., Bach, F. C., Tekari, A., Sagawa, N., Nakamura, Y., Chan, S. C. W., Nakai, T., Creemers, L. B., Frauchiger, D. A., May, R. D., Grad, S., Watanabe, M., Tryfonidou, M. A., & Gantenbein, B. (2018). Successful fishing for nucleus pulposus progenitor cells of the intervertebral disc across species. JOR Spine, 1(2), e1018. 10.1002/jsp2.1018

Salzer, E., Mouser, V. H. M., Tryfonidou, M. A., & Ito, K. (2022). A bovine nucleus pulposus explant culture model. Journal of Orthopaedic Research, 40(9), 2089–2102. 10.1002/jor.25226

Silagi, E. S., Shapiro, I. M., & Risbud, M. V. (2018). Glycosaminoglycan Synthesis in the Nucleus Pulposus: Dysregulation and the Pathogenesis of Disc Disease. Matrix Biology : Journal of the International Society for Matrix Biology, 71–72, 368–379. 10.1016/j.matbio.2018.02.025

Snuggs, J., Basatvat, S., Kanelis, E., Binch, A., Alexopoulos, L., Tryfonidou, M. A., & Le Maitre, C. (2025). Understanding the Physiological Behavior of Disc Cells in an In Vitro Imitation of the Healthy and Degenerated Disc Niche. JOR Spine, 8(4), e70153. 10.1002/jsp2.70153

Snuggs, J. W., Bunning, R. A., & Le Maitre, C. L. (2021). Osmotic adaptation of nucleus pulposus cells: The role of aquaporin 1, aquaporin 4 and transient receptor potential vanilloid 4. European Cells & Materials, 41, 121–141. 10.22203/eCM.v041a09

Spillekom, S., Smolders, L. A., Grinwis, G. C. M., Arkesteijn, I. T. M., Ito, K., Meij, B. P., & Tryfonidou, M. A. (2014). Increased Osmolarity and Cell Clustering Preserve Canine Notochordal Cell Phenotype in Culture. Tissue Engineering Part C: Methods, 20(8), 652–662. 10.1089/ten.tec.2013.0479

Thompson, K., Moore, S., Tang, S., Wiet, M., & Purmessur, D. (2018). The chondrodystrophic dog: A clinically relevant intermediate-sized animal model for the study of intervertebral disc-associated spinal pain. JOR Spine, 1(1), e1011. 10.1002/jsp2.1011

Urban, J. P. G. (2002). The role of the physicochemical environment in determining disc cell behaviour. Biochemical Society Transactions, 30(Pt 6), 858–864. 10.1042/bst0300858

van Dijk, B., Potier, E., & Ito, K. (2011). Culturing Bovine Nucleus Pulposus Explants by Balancing Medium Osmolarity. Tissue Engineering Part C: Methods, 17(11), 1089–1096. 10.1089/ten.tec.2011.0215

van Maanen, J. C., Bach, F. C., Braun, T. S., Giovanazzi, A., van Balkom, B. W. M., Templin, M., Wauben, M. H. M., & Tryfonidou, M. A. (2023). A Combined Western and Bead-Based Multiplex Platform to Characterize Extracellular Vesicles. Tissue Engineering. Part C, Methods, 29(11), 493–504. 10.1089/ten.TEC.2023.0056

van Maanen, J. C., Bach, F. C., Snuggs, J. W., Ito, K., Wauben, M. H. M., Le Maitre, C. L., & Tryfonidou, M. A. (2025). Explorative Study of Modulatory Effects of Notochordal Cell-Derived Extracellular Vesicles on the IL-1β-Induced Catabolic Cascade in Nucleus Pulposus Cell Pellets and Explants. JOR SPINE, 8(1), e70043. 10.1002/jsp2.70043

Vo, N. V., Sowa, G. A., Kang, J. D., Seidel, C., & Studer, R. K. (2010). Prostaglandin E2 and Prostaglandin F2α Differentially Modulate Matrix Metabolism of Human Nucleus Pulposus Cells. Journal of Orthopaedic Research : Official Publication of the Orthopaedic Research Society, 28(10), 1259–1266. 10.1002/jor.21157

Wei, Q., Zhang, X., Zhou, C., Ren, Q., & Zhang, Y. (2019). Roles of large aggregating proteoglycans in human intervertebral disc degeneration. Connective Tissue Research, 60(3), 209–218. (135961823). 10.1080/03008207.2018.1499731

Welsh, J. A., Goberdhan, D. C. I., O’Driscoll, L., Buzas, E. I., Blenkiron, C., Bussolati, B., Cai, H., Di Vizio, D., Driedonks, T. A. P., Erdbrügger, U., Falcon-Perez, J. M., Fu, Q.-L., Hill, A. F., Lenassi, M., Lim, S. K., Mahoney, M. G., Mohanty, S., Möller, A., Nieuwland, R., … Witwer, K. W. (2024). Minimal information for studies of extracellular vesicles (MISEV2023): From basic to advanced approaches. Journal of Extracellular Vesicles, 13(2), e12404. 10.1002/jev2.12404

Williams, R. J., Laagland, L. T., Bach, F. C., Ward, L., Chan, W., Tam, V., Medzikovic, A., Basatvat, S., Paillat, L., Vedrenne, N., Snuggs, J. W., Poramba-Liyanage, D. W., Hoyland, J. A., Chan, D., Camus, A., Richardson, S. M., Tryfonidou, M. A., & Le Maitre, C. L. (2023). Recommendations for intervertebral disc notochordal cell investigation: From isolation to characterization. JOR Spine, 6(3), e1272. 10.1002/jsp2.1272

