## Supplementary Figures 1-4 for "Degenerated intervertebral disc environment impairs notochordal cell-derived extracellular vesicles release and their matrix anabolic effect"

### This document contains:

- Supplementary Table 1
- Supplementary Figures 1-4

**Supplementary table 1.** Details of antibody and protocol for immunohistochemistry on Pig NP explant and Dog NPC pellet.

| **Marker** | **Antigen retrieval** | **Blocking** | **1st antibody** | **2nd antibody** |
|---|---|---|---|---|
| Brachyury | *Pig NP explant & dog NPC pellet*: 10 mM citrate buffer pH 6 (30 min, 70°C) | 0.3% H2O2 (10 min) & mouse serum (ImmunoCruz® goat LSAB Staining System, sc-2053) (30 min) | TBXT (R&D systems, AF2085)<br>*Pig NP explant:* 4 µg/mL<br>*Dog NPC pellet*: 4 µg/mL | Anti-goat biotin/avidin-HRP (Santa Cruz, ImmunoCruz® goat LSAB Staining System, sc-2053) |
| Cytokeratin 8+18+19 | *Pig NP explant*: NA;<br>*Dog NPC pellet*: 10 mM citrate buffer pH 6 (30 min, 70°C) | 0.3% H2O2 (10 min) & 5% PBS/BSA (30 min) | KRT8+18+19 (Abcam, ab41825)<br>*Pig NP explant*: 1 µg/mL<br>*Dog NPC pellet*: 1 µg/mL | Anti-mouse poly-HRP (Immunologic, DPVM110HRP) |
| Angiopoietin-1 receptor | *Dog NPC pellet*: 10 mM citrate buffer pH 6 (60 min, 70°C) | 0.3% H2O2 (10 min) & 5% PBS/BSA (30 min) | TIE2 (Santa cruz, sc-324)<br>*Dog NPC pellet*: 1 µg/mL | Anti-rabbit poly-HRP (Immunologic, DPVR110HRP) |
| Aggrecan | *Pig NP explant*: TE buffer pH 9 (30 min, 70°C) + 10 mg/mL hyaluronidase (45 min, 37°C)<br>*Dog NPC pellet:* 1 mg/mL pronase (60 min, 37°C) + 10 mg/mL hyaluronidase (60 min, 37°C) | 0.3% H2O2 (10 min) & 5% PBS/BSA (30 min) | ACAN (Abcam, ab3778)<br>*Pig NP explant*: 20 µg/mL<br>Dog NPC pellet: 9.67 µg/mL | Anti-mouse poly-HRP (Immunologic, DPVM110HRP) |
| Collagen I | 1 mg/mL pronase (30 min, 37°C) + 10 mg/mL hyaluronidase (30 min, 37°C) | 0.3% H2O2 (10 min) & 5% PBS/BSA (30 min) | COL1 (Abcam, ab6308)<br>*Pig NP explant*: 0.1 µg/mL<br>*Dog NPC pellet*: 0.07 µg/mL | Anti-mouse poly-HRP (Immunologic, DPVM110HRP) |
| Collagen II | 1 mg/mL pronase (30 min, 37°C) + 10 mg/mL hyaluronidase (30 min, 37°C) | 0.3% H2O2 (10 min) & 5% PBS/BSA (30 min) | COL2 (DSHB, II-II6B3)<br>*Pig NP explant*: 0.375 µg/mL<br>*Dog NPC pellet*: 0.03 µg/mL | Anti-mouse poly-HRP (Immunologic, DPVM110HRP) |

**Legend**: NA = not applicable.

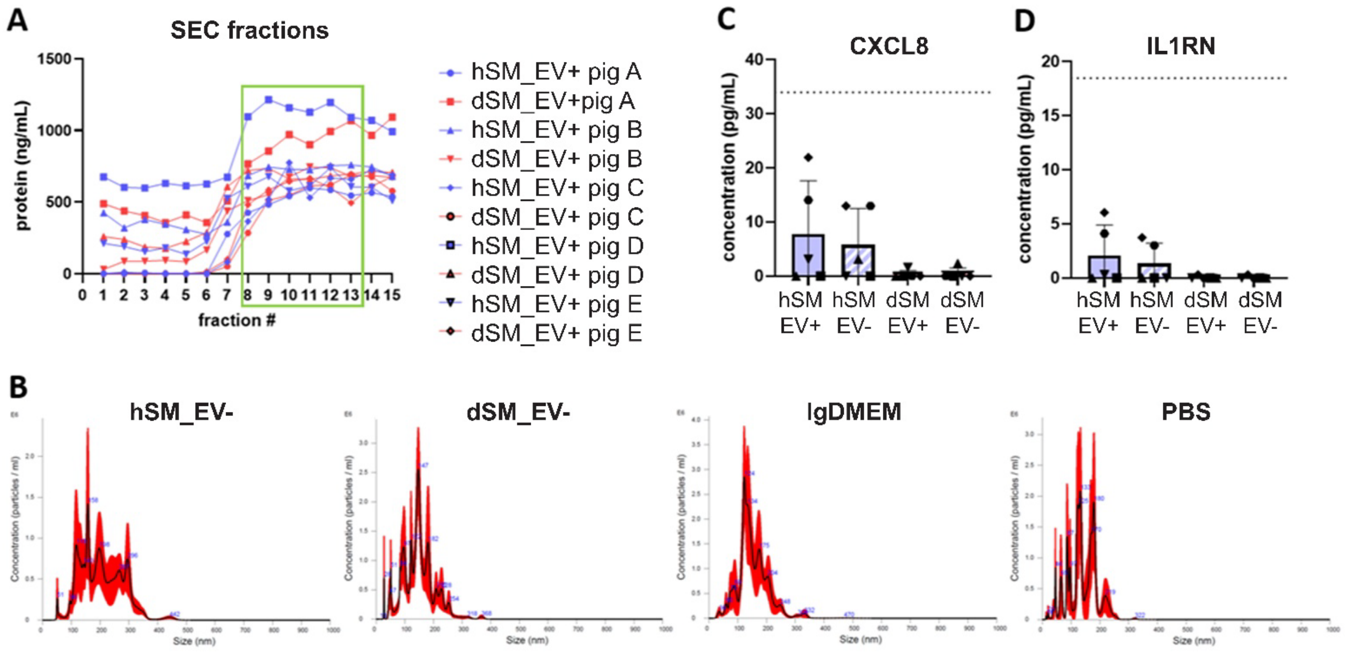


**Supplementary Figure 1. Characterization of healthy and degenerate EV-enriched secretome media (SM_EV+) and respective EV-depleted media (SM_EV-)**

All analyses were done on EV media derived from the conditioned media of pig nucleus pulposus explants cultured in either healthy (hSM_EV+) or degenerate medium (dSM_EV+) for 4 days and their respective EV-depleted controls (hSM_EV- and gdSM_EV-). (**A**) Protein concentration of SEC fractions per pig donor and condition. Green rectangle shows that pooled EV-enriched fractions. (**B**) NTA particle size and concentration profiles of hSM_EV-, dSM_EV-, low glucose medium and PBS. (**C-D**) Levels of CXCL8 and IL1RN in hSM_EV+, hSM_EV-, dSM_EV+ and dSM_EV- media. The dotted line represents the detection limit of respective analytes (CXCL8: 33.97 pg/mL; IL1RN: 18.48 pg/mL). Dots indicate individual matched Pig donors (N=5). Bars and whiskers indicate means ± SD.

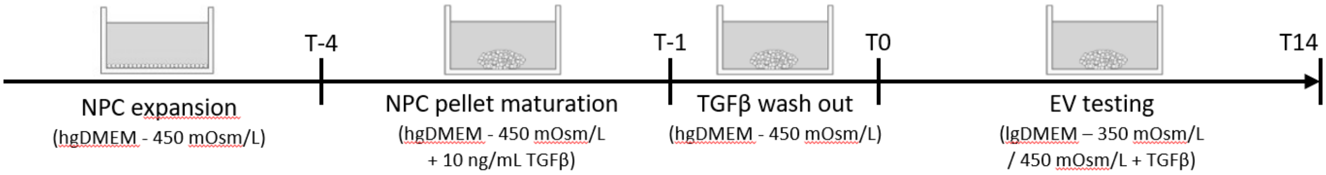


**Supplementary Figure 2. Nucleus pulposus cells (NPC) in vitro 3D model set up.**
Workflow for pellet model production and validation for its capacity to survive and deposit matrix upon TGFβ stimulation either in standard, healthy or degenerative medium.

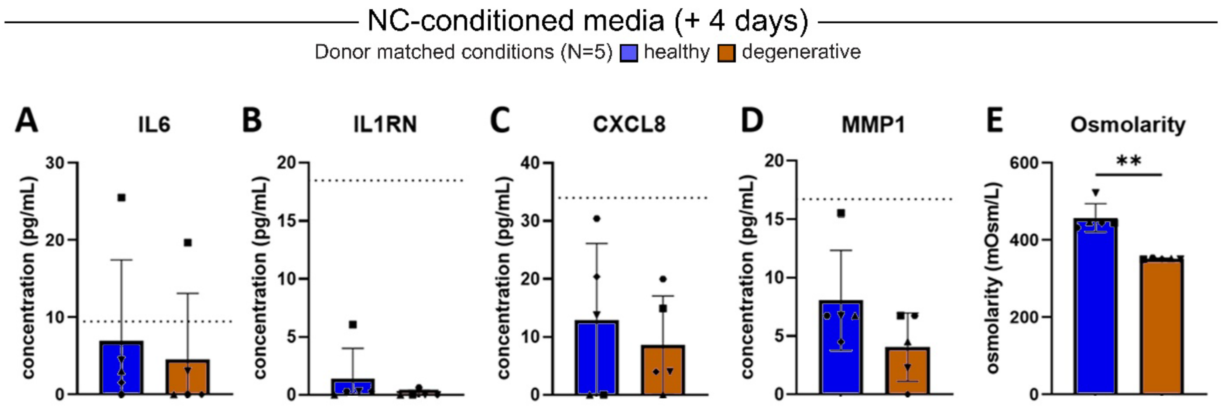


**Supplementary Figure 3. Effect of degenerate media on inflammation factor levels released from NP explants.**

All analyses were done on notochordal cell conditioned media (NCCM) derived from pig nucleus pulposus (NP) explants cultured in healthy or degenerate medium for 4 days. (**A-D**) Levels of IL6, IL1RN, CXCL8 and MMP1 in He- and De-NCCM. Dotted line represents the detection limit of respective analytes (IL6: 9.47 pg/mL; IL1RN: 18.48 pg/mL; CXCL8: 33.97 pg/mL; MMP1: 16.71 pg/mL; INFG: 1.9 pg/mL; IL1B: 2.88 pg/mL; TNF: 12.82 pg/mL). (**E**) Osmolarity measured of NCCM upon 4 days of NP culture – osmolarity of NP culture medium set at day 0 was 450 mOsm for healthy medium and 350 mOsm for degenerate medium. Dots indicate individual matched pig donors (N=5). Bars and whiskers indicate means ± SD. p-values are indicated for statistical significance of differences between conditions using Student's t test (** : $p < 0.005$).

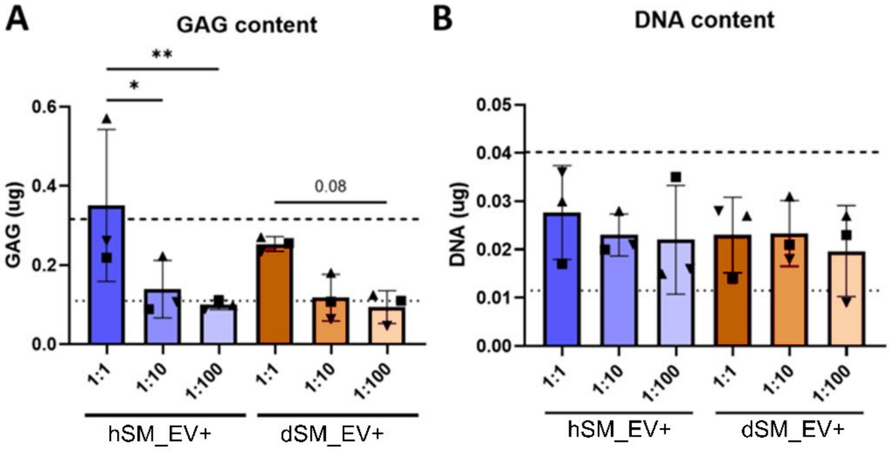


**Supplementary Figure 4**. **Effect of serial dilution of porcine EV-enriched secretome media (SM_EV+) on dog nucleus pulposus cells (NPCs)**

NPC pellets were cultured for 14 days in either degenerate medium (negative control), healthy medium supplemented with 10 ng/mL TGFβ (positive control), or degenerate medium supplemented with SM_EV+ derived in healthy (hSM_EV+) or degenerate (dSM_EV+) medium. (**A-B**) GAG and DNA content in NPC pellet. Dots indicate individual matched pig donor-derived NC-EV treatment (N=3). Bars and whiskers indicate means ± SD. Dotted line and bold dotted line represent negative and positive conditions, respectively. *p*-values are indicated for statistical significance of differences between conditions using Sidak's multiple comparison test (numeric *p*-value: $p < 0.1$, * : $p < 0.05$, ** : $p < 0.005$). Statistical trend $p = 0.05 – 0.1$ is provided in absolute *p* values.
